# spaTransfer: transfer learning for single-cell and spatial transcriptomics data using non-negative matrix factorization

**DOI:** 10.64898/2025.12.12.694021

**Authors:** Cindy Fang, Kelsey D. Montgomery, Sarah E. Maguire, Anthony D. Ramnauth, Boyi Guo, Ryan Miller, Joel E. Kleinman, Thomas M. Hyde, Keri Martinowich, Kristen R. Maynard, Stephanie C. Page, Stephanie C. Hicks

## Abstract

Recent advances in spatially-resolved transcriptomics have enabled profiling of gene expression in a spatial context, which has led to the generation of large-scale single-cell and spatial atlases with computationally-derived cell type or spatial domain labels. An increasingly important task with these data has become the transfer of cell type or spatial domain annotations from a given reference (or source) atlas into a new target tissue or sample. The reference and target datasets could be at different resolutions or measured on different experimental platforms. Here, we present a method to perform cross-platform transfer learning that takes as input single-cell or spatial domain labels from a reference atlas or dataset and transfers the labels to a target dataset at a similar or different resolution. Specifically, we use non-negative matrix factorization (NMF) on the reference data to identify factors associated with labels of interest and project these factors into the target dataset to label each new observation. We use a multinomial model with the factors as covariates and labels as the response to predict labels in the target dataset. In contrast to existing approaches, the advantage of our approach is interpretability, without compromising on accuracy. We demonstrate the performance of the method in two human brain tissues and show that our model identifies spatially coherent domains in the target datasets with concordance of marker gene expression. We implement spaTransfer in open-source software as an R package (github.com/cindyfang70/spaTransfer).

## 1 Introduction

The advent of spatially-resolved transcriptomics (SRT) has led to the generation of large-scale SRT atlases with labeled cell types or spatial domains [1–4]. As new SRT datasets are generated, an important task is to annotate the datasets [5, 6]. In single-cell/nucleus RNA-sequencing (sc/snRNA-seq) data, computational methods have been developed to transfer labels from a sc/snRNA-seq reference atlas to a target scRNA-seq [7–12] or scATAC-seq [13] dataset. These are broadly referred to as “single-cell reference mapping” or “label transfer” algorithms where the goal is to align the -omic (e.g. transcriptomic) profiles of samples in a target dataset to a reference atlas in order to transfer annotations at a single-cell resolution to the target dataset. Recent work has described how the *interpretability* of single-cell reference mapping algorithms is important, especially in applications where the goal is to label complex populations of cells, for example, in response to perturbations [12]. Furthermore, the work suggests computational methods to decompose differential sources of variation and to provide interpretability of the reference mapping algorithms will be beneficial to address reference mapping challenges.

One goal of reference mapping with SRT data is to transfer the labels (either cell types or spatial domains) in the reference atlas to a target dataset. Existing single-cell reference mapping algorithms such as singleR [14] and scmap [15] rely on measures of correlation between gene expression profiles of cells, but imaging-based SRT are limited in the number of measurable features and calculating the correlation between a target and a reference expression profile with different sets of features is challenging. Another challenge is the lack of an ideal molecular atlas due to rapidly emerging new SRT technologies that vary in their resolution. Current approaches to address this challenge include integrative methods such as Cell2location [16] or RCTD [17], which require users to provide paired sc/snRNA-seq reference samples as input. However, these approaches are better suited for decomposing the proportion of cell types rather than transferring cell type or spatial domain labels, as the sc/snRNA-seq reference does not contain any spatial information. Another broad category of integrative label transfer algorithms, such as Symphony [9], are algorithms that learn a low-dimensional representation of the reference dataset and use either cell or gene anchors to map the target dataset into the same low-dimensional space. Unfortunately, many of these algorithms rely on principal component analysis (PCA) for low-dimensional embeddings [12], which yields principal components (PCs) that can only be interpreted sequentially and not as standalone factors. Finally, recent methods have been developed to transfer labels from single-cell reference atlases to SRT data, but these approaches either are designed for similar sets of features or similar experimental platforms, reducing their impact as reference atlases grow across technologies or use deep learning [18–22], which limits their interpretability.

To improve compatibility with diverse SRT technologies across throughput and spatial resolution as well as interpretability to inform biological mechanisms, we developed a novel computational method to transfer labels (cell types or spatial domains) from a reference dataset to a target dataset using non-negative matrix factorization (NMF). This method offers unique advantages to address the challenges noted above by decomposing the observed gene expression into latent factors representing cell types or spatial domains, thus circumventing direct comparisons of expression profiles from differing gene panels. As these factors can be associated with biological proceses, our approach provides direct interpretability of the annotations in the target dataset. In the next two sections, we provide a detailed overview of the method (spaTransfer) and describe the key innovations and advantages it has over existing approaches. Finally, we demonstrate the performance of spaTransfer on several real-world datasets and show how our approach offers clear interpretability without a loss in accuracy compared to existing methods.

## 2 Results

### 2.1 spaTransfer enables cross-platform transfer learning

spaTransfer is a reference mapping algorithm that transfers labeled spatial domains or cell types from a reference (or source) dataset to an unlabeled target dataset, where the datasets can be from different technologies, such as sn/scRNA-seq or SRT platform. Broadly, our approach begins by using non-negative matrix factorization (NMF) to identify factors (or patterns) in the reference dataset (Figure 1). The number of factors (rank), *L* chosen is either a user-specified number or is automatically determined [23]. NMF provides a way to factorize the gene expression data from single-cell or SRT experiments into two matrices with non-negative entries. Briefly, NMF decomposes a matrix 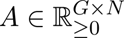, typically gene expression with *G* genes and *N* cells, into two matrices *W* and *H*, such that

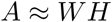

where 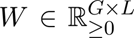 is a gene by rank matrix and 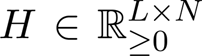 is a rank by cell matrix. For a given target dataset *A*^′^, we solve for *H*^′^ in the equation *A*^′^ = *WH*^′^, using the loadings matrix *W* from the reference dataset. In the setting where the reference dataset and target dataset differ in the observed genes, we subset *W* to only the genes shared between the two datasets. This yields a set of factors in the target dataset that correspond to the factors computed in the reference dataset, which represents the same biological processes, such as cell types or spatial domains. The reference factors (*H*^′^) are then used as features in a multinomial model: regressing the reference factors to the spatial domain labels using the reference dataset. Leveraging the fitted multinomial model, the label for each observation can be predicted using the target dataset. Since it is possible for factors to represent platform-specific or sample-specific factors that are not desirable for downstream prediction, regularization is built into the multinomial model fitting process through elastic net regularization [24, 25].

**Figure 1:**
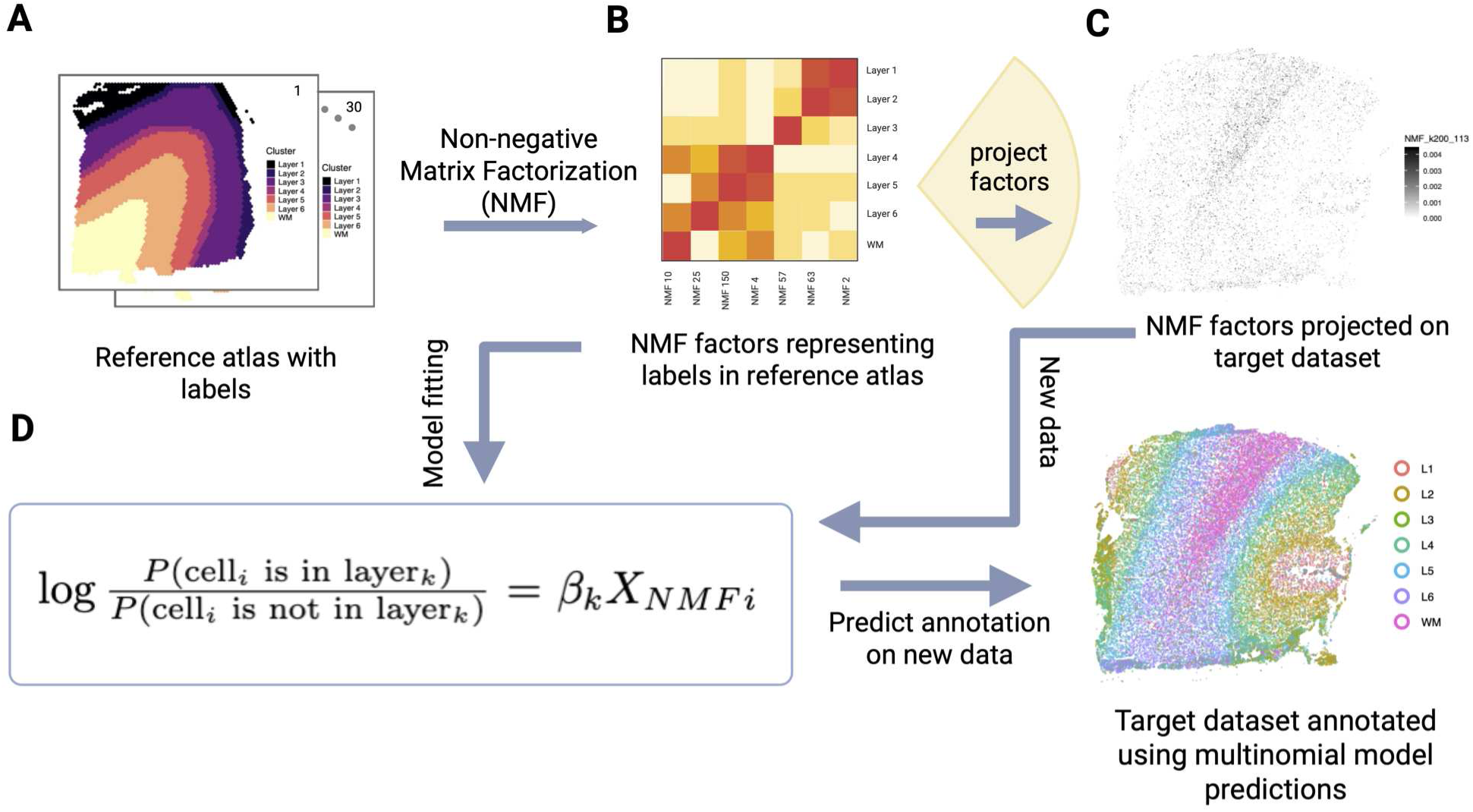
Overview of spaTransfer for transfer learning with single-cell and spatial transcriptomics data using non-negative matrix factorization. (A) Given a set of reference (or source) atlas with multiple samples and labeled cell types or spatial domains, the spaTransfer algorithm uses non-negative matrix factorization (NMF) on the source data to (B) identify factors associated with labels of interest. (C) Next, spaTransfer projects these factors onto the unlabeled target dataset. (D) To label each new observation (e.g. cell or spot), we use a multinomial model with the factors as covariates and labels as the response to predict labels in the target dataset. The primary advantage of our approach is interpretability without compromising accuracy of the label transfer. We demonstrate how our model identifies spatially coherent domains across multiple target datasets.

### 2.2 Overview and key features spaTransfer

spaTransfer enables the transfer of labels between datasets generated from different single-cell or SRT technologies with potentially different sets of features or at different resolutions. For example, spot-based data such as Visium can be thought of as “mini-bulk” data, with expression from multiple cells aggregated into each spot whereas imaging-based data such as Xenium is at single cell resolution. Despite the differing resolutions, there is evidence that NMF factors can address cross-platform challenges. For example, previous studies have shown that NMF factors learned on bulk RNA sequencing data can be transferred to scRNA-seq data, allowing for shared biological features that were first identified in the bulk data to be found in the single cell level data [26]. We take advantage of these properties to transfer information across different technologies. Specifically, we learn the latent factors that can be used to approximately generate the reference dataset through additive linear combinations. spaTransfer then estimates the composition of these latent factors in the target dataset. Based on the distribution of latent factors in each observation, spaTransfer is able to predict the annotation for the observation. Previous SRT studies have leveraged these latent factors for both discrete cell types and continuous patterns of gene expression [27, 28]. Overall, spaTransfer provides a unified framework for performing NMF on the reference dataset, transferring NMF factors to the target dataset, and leveraging these factors to fit predictive label transfer models.

Second, spaTransfer offers a computationally fast and biologically interpretable way to label cells or spatial domains in an unlabeled target dataset. Previously, NMF was challenging to use with high-dimensional transcriptomic datasets because of the computational burden to compute the factors. However, recent advances in optimization for matrix factorization for sparse matrices [23] have led to implementations of NMF that are comparable in computational speed to well-established algorithms that compute partial singular value decompositions (SVD) of large sparse or dense matrices [29, 30]. This greatly reduces the computational burden of using NMF in practice to find low-dimensional representations of transcriptomics data, such as for transfer learning. Furthermore, the additive nature of the algorithm enables us to sift out signatures representing interesting biological patterns, with each factor representing distinct biological processes[23]. The non-negative nature of NMF is particularly suited to transcriptomics data, as gene expression counts are inherently non-negative. This strategy allows for a more interpretable transfer learning algorithm than relying on PCA, which can result in negative values. NMF on the atlas yields a ‘loadings matrix’, which will be used to compute projections of the factors in the new dataset. These projected factors can then be used to identify tissue areas or cells with enrichment of biological patterns.

Third, spaTransfer can leverage information from multiple reference samples due to its usage of NMF. The additive nature of NMF allows for (i) each factor to be interpreted independently and (ii) removing factors that are highly correlated with unwanted technical artifacts, thus reducing the impact of potential erroneously labeled observations in the target dataset due to batch effects observed in the reference dataset. Fourth, spaTransfer offers the option to smooth predicted labels in the target dataset across space. This step is especially helpful when transferring spatial domain labels as it increases the spatial contiguity of the predicted domains. Spatial smoothing of domains controlled by density of labels was also presented in BANKSY [31], but spaTransfer’s smoothing algorithm allows for domain-specific density parameters rather than a single global parameter.

### 2.3 spaTransfer provides interpretable factors to transfer labels from a reference atlas to a target dataset across experimental platforms with different resolutions

Next, we consider two examples where we apply spaTransfer and show how our model transfers spatial domain labels obtained in a reference dataset to an unlabeled target dataset at a single-cell resolution. The purpose is to demonstrate how spaTransfer is able to identify spatially coherent domains in the target dataset, which is profiled using a different experimental platform at a different resolution compared to the reference dataset.

#### 2.3.1 Application to human dorsolateral prefrontal cortex

The granular cytoarchitecture of the human dorsolateral prefrontal cortex (dlPFC) is linked to the region’s key role in higher-order cognitive functioning. Previous studies profiled human dlPFC tissue using snRNA-seq (single cell resolution) and the 10x Genomics Visium platform based on spots (multiple cells / spot resolution) [32–38]. The dlPFC contains six classic histological layers, which each have different distributions and densities of distinct cell types. To investigate the distribution of cell types within individual spatial domains (e.g. corresponding to histological layers), we generated single-cell resolution 10x Genomics Xenium data in postmortem human dlPFC tissue (6 tissue sections across 4 neurotypical brain donors) using the Human Brain Gene Expression panel (266 unique genes) (Figure 2A, Table S1, Methods). Next, we leveraged a reference dataset with labeled spatial domains measured on the 10x Genomics Visium platform with 30 tissue sections across 10 neurotypical brain donors [33] (Figure 2B).

**Figure 2:**
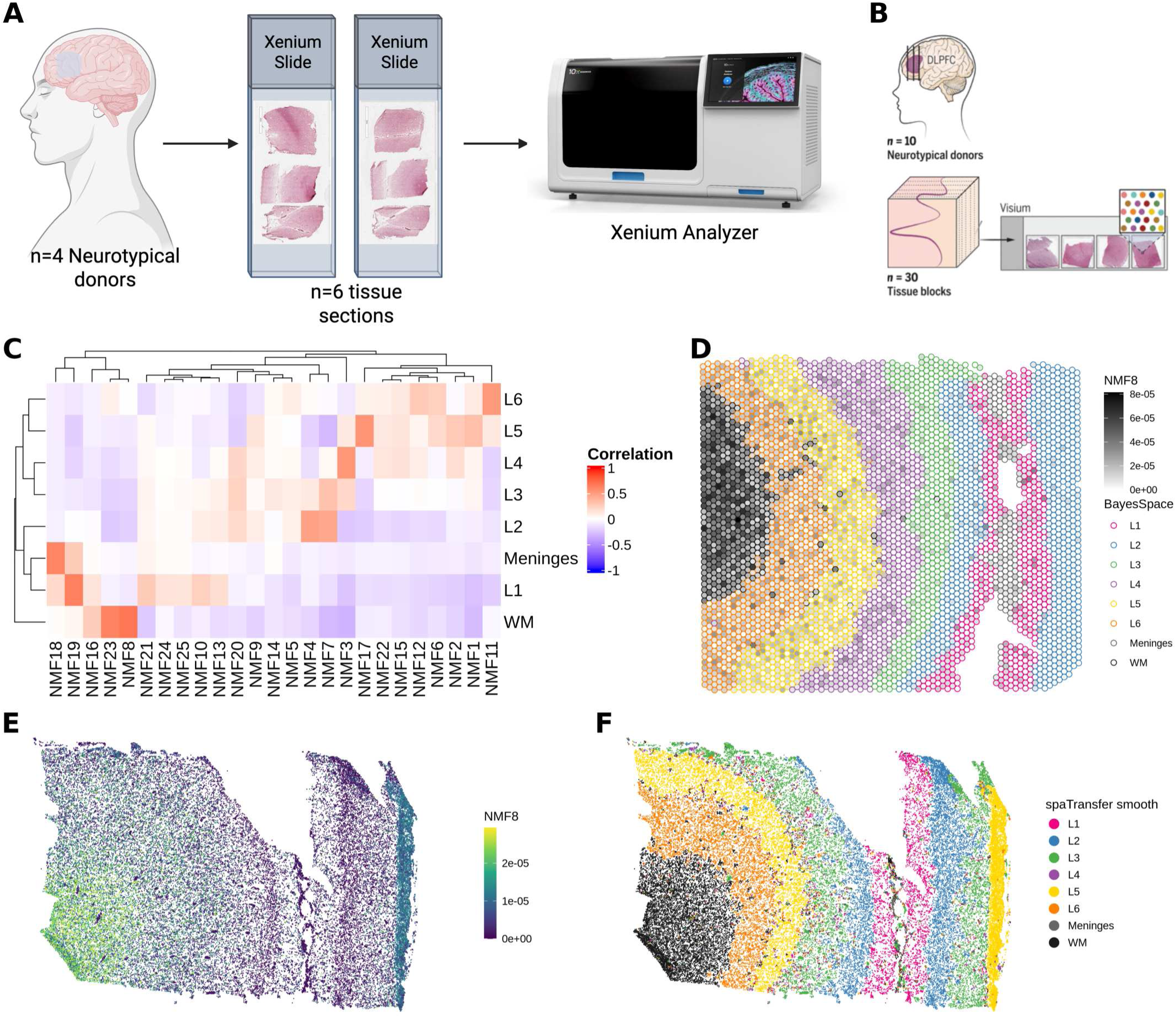
spaTransfer accurately transfers spatial domain annotations to a target dataset at single-cell resolution in human dorsolateral prefrontal cortex. We used two datasets both generated in postmortem human tissue from dorsolateral prefrontal cortex (dlPFC): (1) (A) an unlabeled target dataset at single-cell resolution measured on the 10x Genomics Xenium platform with 6 tissue sections generated across 4 neurotypical donors, and (2) (B) a reference dataset with labeled spatial domains measured on the 10x Genomics Visium platform with 30 tissue sections across 10 neurotypical donors [33]. (C) Heatmap of factors learned from the reference dataset using spaTransfer, which highly correlate with labeled spatial domains. (D) Spotplot of weights for factor 8 (NMF8) that is associated with white matter (WM) and visualized with labeled spatial domains in a tissue sample (‘Br6471 Post’). (E) Cell-level plot of NMF8 projected in a Xenium tissue sample (‘Br6471_Post_5434’). (F) Same tissue as E, but cell-level plot with colors representing the predicted spatial domains.

Using the reference Visium dataset, we used spaTransfer to transfer the spatial domain labels to the unlabeled target dataset measured on the Xenium platform. First, we applied standard preprocessing and quality control (QC) considering negative control genes (Figures S1-S3) in the target dataset. Next, we identified factors using NMF in the reference dataset that were strongly correlated with the labeled spatial domains (Figure 2C) and other unwanted technical factors (Figure S4). For example, we found that factor 21 (NMF21) was associated with spots with high percentages of mitochondrial counts and factor 8 (NMF8) was highly correlated with the white matter (WM) domain in the reference dataset. This is an example of the interpretability provided by spaTransfer. Then, we visualized the factor loadings for NMF8 in 2D resolution on a Visium tissue section, which showed the factor weights were localized to the regions annotated as WM (Figures 2D). These factors were then projected into the Xenium dlPFC target datasets. The projected factors retained their spatial organization and aligned with known anatomy (Figures 2E, S5). Specifically, we found that spatial domain predictions from spaTransfer based on these transferred factors matched the expected laminar structure of the dlPFC, with individual cortical layers correctly annotated and sequentially positioned from the more superior meninges to the deeper WM layers (Figures 2F, S6, S7). Interestingly, some Xenium tissue slices such as those from donor Br8667 did not contain WM. This information was not provided to the method for label transfer, but spaTransfer was still able to accurately identify the structure of the tissue slice (Figures S6, S7).

#### 2.3.2 Application to human dentate gyrus (DG)

Structural and functional synaptic plasticity in neurons of the hippocampus (HPC), particularly those within the dentate gyrus (DG) subfield, underlie several forms of learning and memory that change dynamically over the lifespan [40–42]. Many functions of the DG are topographically organized, and therefore, a better understanding of gene expression patterns across DG cell types, and how they change over development and aging, is important [39, 40, 43]. The human HPC and DG has been profiled using snRNA-seq across the lifespan, but this approach does not retain spatial information [44–48]. We recently used SRT to profile postmortem human DG tissue sections from 16 donors across the lifespan (N=5 infant, N=4 teen, N=4 adult, N=3 elderly) using 10x Genomics Visium. Our study identified heterogeneity in spatial gene expression within the DG for molecular markers of proliferation, extracellular matrix, and neuroinflammation [39]. However, information from individual cells is not available on the Visium platform.

To facilitate better understanding of the human DG by utilizing both spatial and single-cell information, we generated SRT data on a subset of the donors from our previous study [39] using a precommercial prototype version of the 10x Genomics Xenium Prime 5K Human Pan Tissue and Pathways Assay. We profiled the postmortem human DG from four brain donors across the lifespan (N=1 infant, N=1 teen, N=1 adult, N=1 elderly) (Figure 3A). Similar to the Xenium platform, Xenium Prime profiles gene expression at a single cell resolution, but allows for up to 5000 genes to be measured. First, we applied standard preprocessing and QC (Figures S8-S10). To annotate spatial domains in this newly generated dataset, we applied spaTransfer using the spatial domain labels from the reference atlas (N=16 Visium DG dataset) and the newly generated Xenium datasets as the target dataset. The Visium DG dataset was previously annotated by performing unsupervised, spatially-aware clustering with BayesSpace [49] followed by identifying canonical marker genes of each cluster [39]. These annotations consist of *k*=10 clusters each representing a unique spatial domain (Figure 3B). spaTransfer was able to identify latent factors representing spatial domains in the Visium reference dataset as quantified by Pearson correlation (Figure 3C). We also examined the spatial localization of these factors by plotting the factor values for each spot on the tissue. These plots showed that the spatial patterns of these factors are aligned with known anatomy (Figure 3D) and the BayesSpace annotations. Projecting these factors into the Xenium DG dataset using spaTransfer yielded factors that exhibited spatial localization concordant with anatomical structures. The spatial localizations of the projected factors were consistent between the reference and target datasets, for example NMF1 represents the granule cell layer (GCL) in the reference dataset (Figure 3D) and the projected NMF1 in the target datasets show spatial localization in the expected location of the GCL (Figures 3E, S11). Lever-aging the NMF factors, which encode both spatial and gene expression information, spaTransfer was able to predict spatial domain annotations for each cell in the Xenium DG dataset. These predicted annotations show concordance with the known anatomy of the region and enable spatially-aware downstream analyses (Figures 3F, S12, S13).

**Figure 3:**
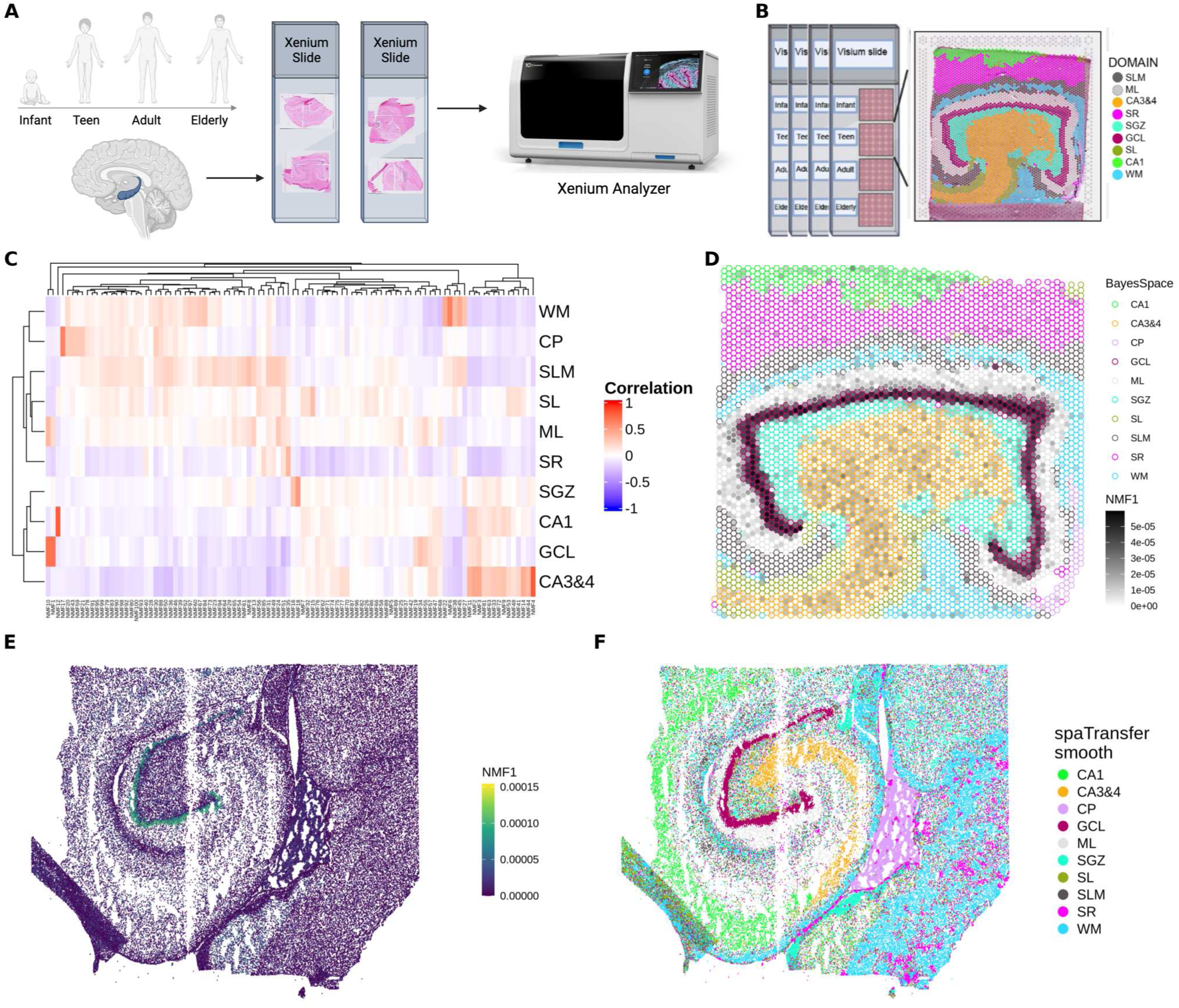
spaTransfer accurately transfers spatial domain annotations to a target dataset at single-cell resolution in postmortem human dentate gyrus (DG) of the hippocampus. We considered the following two datasets: (A) an unlabeled target dataset at a single-cell resolution measured on the 10x Genomics Xenium Prime platform with 4 tissue sections of human dentate gyrus from 4 neurotypical brain donors. Each donor represented one stage across the human lifespan (infant, teen, adult, elderly). (B) A reference dataset with labeled spatial domains measured on the 10x Genomics Visium platform, with 16 tissue sections from 16 donors, profiling the DG across the lifespan [39]. (C) Heatmap of factors learned from the reference dataset using spaTransfer, which highly correlate with labeled spatial domains. (D) Spotplot of weights for factor 1 (NMF1), which is associated with the granule cell layer (GCL) visualized with labeled spatial domains in tissue sample Br1412_teen (E) Cell-level plot of NMF1 projected in the Xenium tissue sample Br8533_infant. (F) Same as (E), but cell-level plot with colors representing the predicted spatial domains.

Finally, we used the transferred labels in the Xenium DG data to explore cell-level differences (annotated with the spatial domains) across the lifespan. Specifically, we pseudobulked the cell-level counts across the gene for each spatial domain in each donor. After removing pseudobulk outliers related to the choroid plexus and other donor-specific artifacts (Figure S14), PCA was applied to the remaining 30 pseudobulk samples to identify the top components of variation in the data. We found that the first principal component (PC) (46% of variance explained) appears to be driven by differences between broad cell classes (neurons, white matter, neuropil) (Figure S15). Separation of infant samples from the other age groups is seen along PC2 (15% of variance explained), which matches findings in a previous study [39]. This suggests that gene expression patterns in the DG during early life significantly differ from that in later life (Figure S16).

### 2.4 spaTransfer accurately annotates spatial domains with high spatial contiguity and cluster validity

To assess the performance of spaTransfer, we considered two sets of evaluations. First, we aimed to evaluate the accuracy of annotating spatial domains using spaTransfer compared to three widely used single-cell reference mapping algorithms, namely singleR [14], Seurat [10, 50], and scmap [15]. This enables the comparison of spaTransfer’s prediction accuracy against other label transfer algorithms. Second, we considered the performance of spaTransfer compared to unsupervised spatial domain detection algorithms to evaluate the spatial contiguity and cluster validity. Our rationale for this was to evaluate the performance of algorithms that transferred labels from a reference dataset compared to algorithms that perform *de novo* identification of spatial domains in a new dataset using unsupervised clustering algorithms. We recognize these can be complementary approaches, but with the increasing availability of reference datasets and atlases, we aimed to answer this question more formally.

Towards the first goal, we considered the human dlPFC data profiled on the 10x Genomics Visium platform [33] as described in Section 2.3.1. Specifically, we split the 30 tissue sections from this dataset in half where 15 were used as the reference dataset and the other 15 were used as the target samples. We used the originally reported spatial domains as the labels in the reference atlas for spaTransfer, singleR [14], Seurat [10, 50], and scmap [15] and transferred the labels using these algorithms (see Methods for details). To compare the accuracy between these methods using the 15 target samples, we used the originally reported data-driven spatial domain labels (using BayesSpace [49]) in the 15 target samples to compare against the transferred labels from the three algorithms (Figure 4A). To assess the accuracy of the label transfer methods, we computed the adjusted Rand index (ARI) and normalized mutual information (NMI) (Figure 4B). We found that spaTransfer (median ARI: 0.49, median NMI: 0.59) outperforms singleR (median ARI: 0.16, median NMI: 0.26), Seurat (median ARI: 0.43, median NMI: 0.50), and scmap (median ARI: 0.27, median NMI: 0.32) in terms of both ARI and NMI, as higher values indicate better performance.

**Figure 4:**
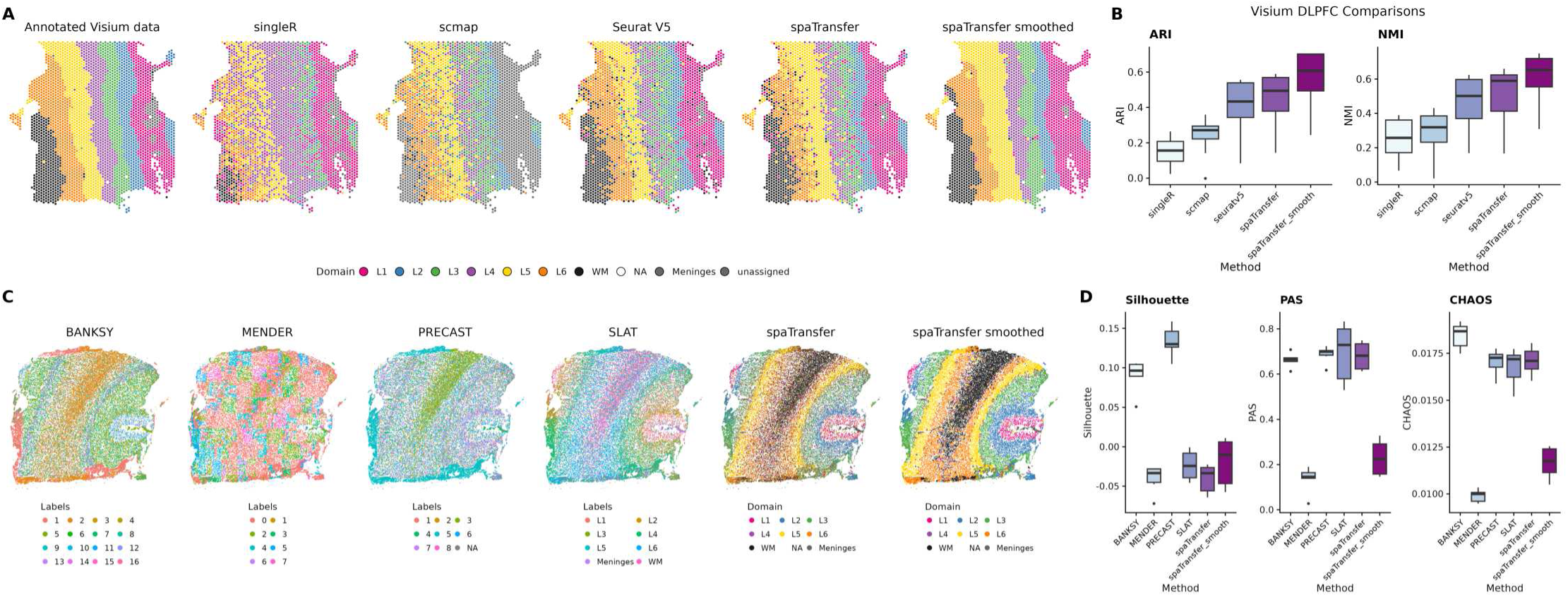
spaTransfer accurately annotates spatial domains with high spatial contiguity and cluster validity. (A) Predictions from all transfer-learning based methods on Visium tissue Br6471_ant. (B) Boxplots of ARI and NMI for each of the different label transfer methods, computed across the target set Visium samples. spaTransfer shows better average performance than singleR in terms of both ARI and NMI. spaTransfer also shows comparable performance to Seurat. (C) Cluster labels from all unsupervised methods on Xenium tissue Br2743_Mid_5548. (D) Silhouette index, proportion of abnormal spots (PAS), and CHAOS metrics computed for all unsupervised methods across all 6 Xenium dlPFC samples. spaTransfer has the best performance in terms of the PAS and CHAOS metrics.

We also evaluated the performance of spaTransfer on Xenium data against widely used spatial domain detection methods that can be applied to imaging-based SRT, namely BANKSY [31], MENDER [51], PRECAST [52], and SLAT [53]. For this evaluation, we continued with the same human dlPFC data profiled on the 10x Genomics Xenium platform as described in Section 2.3.1. To evaluate the biological validity of the predicted domains, we used an internal validity metric that did not depend on a ground truth label, namely the silhouette value [54], which was calculated on the top 50 PCs of normalized gene expression. The maximum silhouette value is 1 and the minimum value is −1, with higher scores indicating better performance. Next, we aimed to evaluate the spatial contiguity of the predicted domains or how “domain-like” each cluster was, with the assumption being that spatial domains should exhibit contiguity in physical space. Towards this goal, we considered two metrics: (i) proportion of abnormal spots (PAS) [55] and (ii) spatial chaos (CHAOS) [55]. For both of these metrics, lower values of the metrics indicate more spatial contiguity. In our evaluations, we found that spaTransfer reported the highest spatial contiguity compared to reference-free methods as quantified by PAS, and is competitive in terms of CHAOS (Figures 4D, S17). Notably, the performance of spaTransfer was lower using the silhouette index in our evaluations. However, the silhouette index was computed using principal components of gene expression, meaning it was inherently biased towards BANKSY, MENDER, and PRECAST, which all rely on PCA for dimension reduction before clustering.

### 2.5 Sensitivity analysis of factorization parameters

Finally, we assessed the sensitivity of spaTransfer’s performance to parameter and design choices. We first sought to quantify the prediction performance of spaTransfer across a range of NMF factorization ranks. To do so, we again used the Visium human dlPFC data described above, using half the samples as the reference and the other half as the target. We then ran spaTransfer at ranks of 5, 10, 15, 20, 25, 40, 60, 100, 150, 200, 300, 400, and 500. At each of the ranks, we quantified the prediction accuracy using ARI and the reconstruction accuracy using mean squared error (MSE) on held-out matrix entries. We observed that ARI began to plateau and MSE start to increase around a rank of 25, indicating that this rank balances prediction performance and reconstruction faithfulness (Figure 5A). Interestingly, we noticed that no single rank optimizes both ARI and MSE, suggesting that choosing an optimal rank should depend on the user’s priorities and downstream applications.

**Figure 5:**
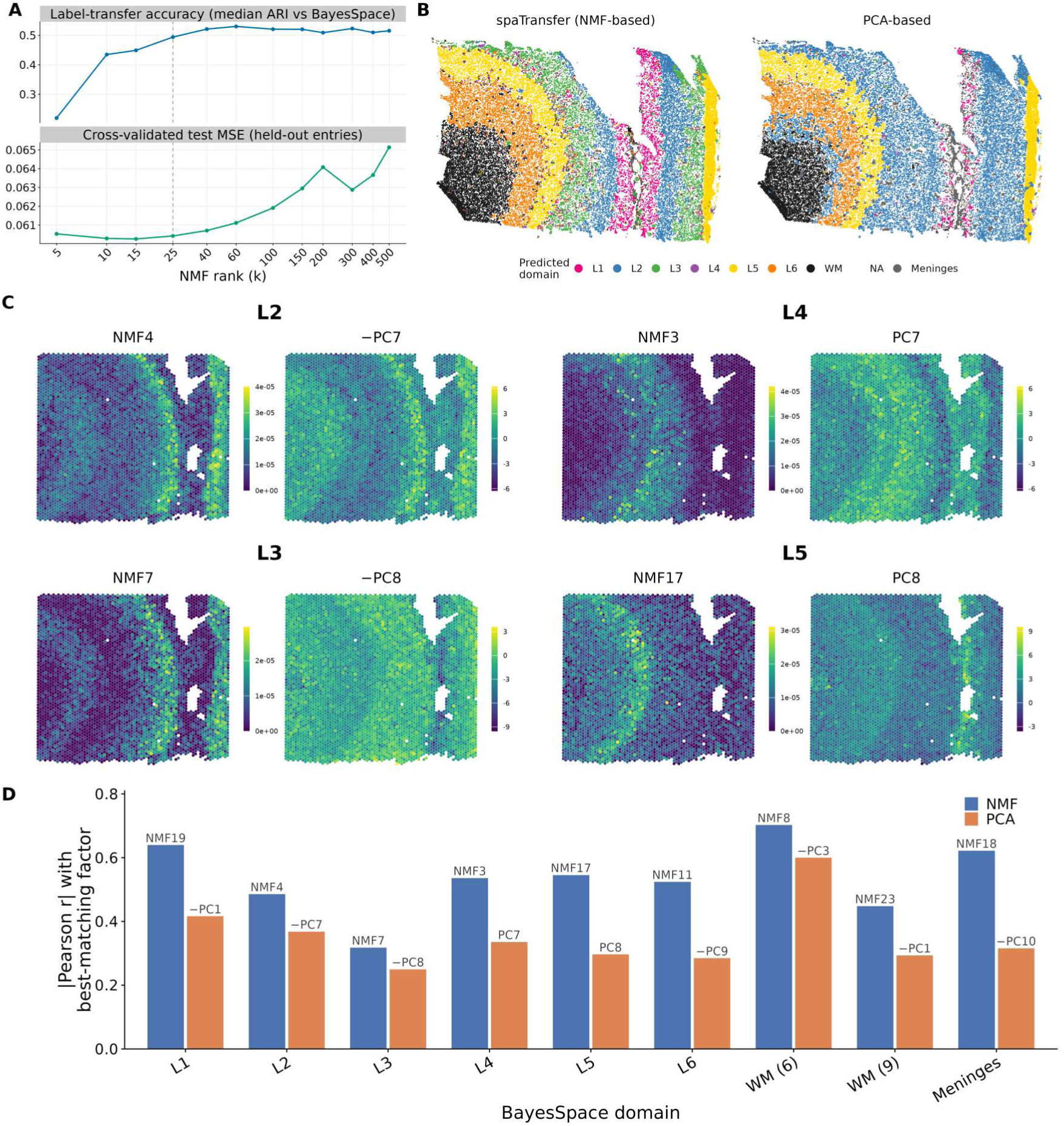
Rank and factorization sensitivity analysis of label transfer. (A) Rank sensitivity analysis of spaTransfer. Top: sensitivity of prediction accuracy across NMF ranks ranging from 5 to 500, quantified using ARI. Prediction accuracy is highest at a rank of 60. (B) Comparison of NMF and PCA for factorization-based prediction performance. Predicted labels on tissue section Br6471_Post_5434 from spaTransfer (left) and a PCA-based workflow (right) using a rank of 25 for both. The PCA-based approach notably lacks clean L1 and L3 domains, and has predicted L2 incorrectly directly to the right of WM. (C) Tissue plots of the NMF and PCA factors with the highest absolute correlation to L2, L4, L3, and L5, respectively. NMF factors appear to be more spatially localized to the expected region of each domain, whereas PCA factors are more diffuse. As well, NMF factors are unique across the domains whereas PC7 is associated with both L2 and L4, and PC8 is associated with both L3 and L5. (D) Quanitfying how well factors from NMF and PCA capture relevant biological information. For each spatial domain, we compared the NMF and PCA factors with the highest absolute correlation with the domain. Across all domains, NMF factors have higher absolute correlation than PCA factors. NMF factors are also more interpretable as they all have positive correlation with biological domains, whereas several PCA factors are negatively correlated with domains.

We next evaluated the choice of NMF for the factorization step in spaTransfer by comparing it to PCA while keeping all other parameters and design choices the same. We repeated the procedure in 2.3.1 for both spaTransfer and the PCA-based workflow, using the same factorization rank for both. We observed that spaTransfer predictions better recapitulated cortical layers as the PCA-based workflow lacked clean L1 and L3 domains, and predicted L2 in areas where it shouldn’t exist (Figures 5B, S19). To better understand the information contained in each of the factorizations, we computed the Pearson correlation between each factor and the ground-truth Visium spatial domains. For each spatial domain, we plotted the NMF and PCA factors with the highest correlation on the tissue section. We observed that NMF factors appeared more spatially localized to the expected region of the spatial domain (Figure 5C) We found that NMF factors were consistently more highly correlated than PCA factors (Figure 5D). NMF factors were all positively correlated with spatial domains, whereas several PCA factors showed negative correlation and were therefore harder to interpret. As well, PCA factors often were associated with multiple domains, for example PC7 had high correlation with L4, but was also anticorrelated with L2. Taken together, spaTransfer’s choice of NMF provides higher prediction accuracy and greater interpretability than PCA-based approaches.

## 3 Discussion

We introduced a new computational method spaTransfer to transfer the labels (cell types or spatial domains) from a reference atlas to an unlabeled target dataset that prioritizes interpretability and can be used across different platforms with different sets of features. The core of our method using non-negative matrix factorization to learn a set of latent factors in the reference atlas and then use a multinomial model to predict the labels (e.g. spatial domains or cell types) using the latent gene expression patterns.

We demonstrate the performance of spaTransfer using two reference atlases generated using the Visium platform [33, 39] to transfer the labels to two newly generated Xenium datasets (human dlPFC and human dentate gyrus). These datasets complement existing Visium datasets by providing higher-resolution spatial measurements, enabling the study of transcriptional heterogeneity at cellular resolution. Most notably, we demonstrated the ability of spaTransfer to capture and transfer gene expression patterns across different platforms with different sets of features, while also balancing biological validity and spatial contiguity as evaluation metrics. In terms of performance, spaTransfer accurately identified spatial domains in these datasets and also produced more spatially contiguous domains, which aligned with expected neuroanatomy.

Towards limitations, we recognize that several deep learning algorithms have already been developed to transfer labels between single-cell reference atlases and spatial target datasets. However, these tools lack transparency and interpretability, which are necessary if there is increased reliance on reference atlases to transfer cell type or spatial domain labels to newly generated target datasets. We also acknowledge that NMF requires a user-defined parameter (*L* or the number of latent factors), which could impact the performance of the label transfer algorithm if not chosen carefully. In sensitivity analyses, we identified that the optimal rank that minimizes mean-squared error of reconstruction does not always correspond to the rank that optimizes prediction performance. This is an important area of future research, which has become more computationally feasible due to advances in sparse matrix factorization algorithms [23]. Finally, we recognize that transferring labels from transcriptome-wide technologies to targeted gene panels will be inherently limited by the gene panel. Although our simulation analyses demonstrated that subsetting a dataset to 25% of its original features preserves the distribution of NMF factors (Figure S18), results will likely depend on how informative each gene or feature is. Thus, it is important to carefully select genes during panel design to consider whether genes that are relevant for label transfer will be present.

## 4 Methods

### 4.1 Datasets

#### 4.1.1 Publicly available datasets using 10x Genomics Visium

##### human dlPFC

Gene expression counts were obtained from the *spatialLIBD* R package [56] as described in Huuki-Myers et al. [33]. This reference dataset contains 3 tissue regions spanning the rostral-caudal axis of the dlPFC from each donor, with a total of 10 neurotypical adult donors. The 10x Genomics Visium Spatial Gene Expression platform was used to profiling the gene expression of postmortem dorsolateral pre-frontal cortex (dlPFC) in human brain across *N* =30 tissue sections, 28,916 genes, and *n*=113,927 spots. The reference data was previously annotated using the spatially-aware data-driven clustering method from BayesSpace [49] and spatially registering BayesSpace clusters to manual spatial domain annotations. These spatial domain annotations consist of 7 different labels: Layers 1 through 6 (L1-6) and white matter (WM).

##### human DG

Gene expression counts were obtained from Zenodo [57] as described in Ramnauth et al. [39]. This reference dataset consists of *N* =16 tissue samples of anterior hippocampus from fresh-frozen postmortem human brains. The dentate gyrus (DG) was dissected from these slabs and mounted on Visium slides, with each capture area containing a unique donor. This resulted in 30,217 genes measured across *n*=68,685 spots. The reference dataset clustered using BayesSpace [49] for a total of *k*=10 clusters which were then annotated with canonical hippocampal spatial domains based on expression of marker genes and comparison with histology from H&E staining.

#### 4.1.2 Postmortem human tissue samples

##### Postmortem human tissue samples

Postmortem human brain tissue from donors of European ancestry were obtained at the time of autopsy following audiotaped witnessed informed consent from legal next-of-kin, through the Maryland Department of Health IRB protocol #12–24, and from the Western Michigan University Homer Stryker MD School of Medicine Department of Pathology, and the Department of Pathology at University of North Dakota School of Medicine and Health Sciences, all under the WCG protocol #20111080. One additional sample was obtained by material transfer agreement from the NIMH Brain Collection Core, under protocol #90-M-0142. Curation, tissue handling, processing, and quality control measures have been detailed previously by Lipska et al. [58]. This study included four HPC samples and four dlPFC samples, and demographic information for all donors is listed in Supplemental Table S1. A small piece of dlPFC or DG from each donor was dissected under the visual guidance of a neuroanatomist with a handheld drill on dry ice. Microdissected dlPFC and DG tissues were stored at −80C until cryosectioning, and anatomical orientation was validated as previously described by Huuki-Myers et al. [33], Ramnauth et al. [39].

#### 4.1.3 Generation of human dlPFC using 10x Genomics Xenium

##### Tissue processing

The dlPFC data shown in this study were collected from 4 adult neurotypical donors (Table S1). Cryosections from dlPFC tissue samples were mounted onto Xenium slides with 3 tissue samples from 3 donors on each slide. We collected adjacent replicates across the 2 Xenium slides for 2 donors. Sectioning specifications for dlPFC samples are listed in the 10x Genomics protocol CG000579 (Rev C).

##### Xenium In Situ Workflow

Fixation and permeabilization steps were performed as described in protocol CG000581 (Rev C, 10x Genomics). Probe hybridization, ligation, amplification, autofluorescence quenching, and nuclei staining was performed as specified in CG000581 (Rev C, 10x Genomics) using the 266 gene human brain panel (Xenium Human Brain Gene Expression Panel, Part No.1000599). All slides were loaded onto the Xenium analyzer at the Single Cell - Transcriptomics Core (SC-TC) at the Johns Hopkins University School of Medicine and regions of interest (ROIs) were selected according to the 10x protocol CG000584 (Rev D).

#### 4.1.4 Generation of human DG using 10x Genomics Xenium prime and multimodal segmentation

##### Tissue processing

The DG data shown in this study came from four donors of differing ages (N=1 infant, N=1 teen, N=1 adult, N=1 elderly) (Table S1). Cryosections from fresh frozen DG tissue samples were mounted onto Xenium slides with two tissue samples from different donors on each slide. The infant and adult samples were mounted onto three slides and the teen and elderly were mounted onto another three slides. Sectioning specifications for these samples are listed in the 10x Genomics protocol CG000579 (Rev C). Slides with frozen tissue sections were shipped on dry ice to 10x Genomics, where the samples were processed in two separate batches.

##### Xenium In Situ Workflow

To perform Xenium in situ analysis (10x Genomics) on DG samples, fixation and permeabilization steps were performed as described in protocol CG000581 (Rev C, 10x Genomics). Probe hybridization, ligation, amplification, autofluorescence quenching, and multimodal cellular segmentation staining was performed by 10x Genomics. The gene panels from both batches contain a 4200 gene base panel and a choice between two high expression panels of 289 genes. Genes in the two high expression panels were compared to differential expressed genes from an existing Visium study of the hippocampus across the lifespan. The panel with more overlap with genes that were differentially expressed in age groups was selected. Unlike the first batch, we were also able to provide custom gene targets in the second batch. This custom panel consisted of 15 genes that are important for expression profiling in the DG. These genes are *SLC17A7*, *SLC17A6*, *CD74*, *DCX*, *PROX1*, *LAMP5*, *TTR*, *FIBCD1*, *NECAB1*, *FN1*, *PPFIA2*, *MPPED1*, *SLC12A5*, *METTL7B*, and *SEMA5A*.

### 4.2 Analysis of Xenium datasets

#### 4.2.1 Xenium Onboard Analysis

After imaging on the Xenium Analyzer instrument, the dlPFC and DG data were processed using the 10x Genomics Xenium On Board Analysis (XOA) pipeline. The XOA pipeline processes the volumes generated across different fields of view (FOVs), fluorescence channels, and probe hybridization and imaging cycles. The dlPFC datasets were processed using XOA version 1.5 and the DG datasets were processed using a prototype version of XOA 3.0.0.0. In both versions of the XOA pipeline, image processing and RNA decoding is performed.

#### 4.2.2 Quality control and preprocessing

Outputs from the XOA pipeline were used as input for quality control and preprocessing of the Xenium data prior to performing downstream analyses. Specifically, the cell_feature_matrix.h5, cells.csv.gz, cell_boundaries.csv.gz, and nucleus boundaries.csv.gz files were read into R as using the *SpatialExperiment* package [59]. The cell_feature_matrix.h5 file contains high-quality transcript counts that are found within cells. High-quality transcripts were identified by the XOA pipeline based on a Phred-style score, which is then calibrated using control probes and codewords. Several types of control probes and codewords are used in the Xenium workflow, and a brief description of each type of negative control is available from 10x Genomics. We refer to these collectively as ‘negative controls’. Only counts with a calibrated score of ≥ 20 are deemed high-quality and retained for downstream analyses. The cell feature matrix is then used as the counts matrix in a SpatialExperiment object. The cells.csv.gz file contains the spatial coordinates of each cell as well as each cell’s area, its nucleus area, the total number of counts in transcripts, the number of counts from negative controls, and the total number of overall counts. Due to the limited gene panel in Xenium, we do not quantify reads mapping to the mitochondrial genome and therefore do not use this as a QC metric. Instead, the *scater* R package [60] was used to identify and remove low quality cells based on thresholding counts from negative controls. Cells that were 5 or more median absolute deviations (MADs) higher than the median value of any negative control were deemed to be low quality and removed from downstream analyses. Additional filtering was performed to remove cells with zero total counts and a nucleus area of less than or equal to 200 microns squared, based on expected cell sizes in the human brain. For the Xenium dlPFC dataset, 17,972 cells were discarded out of a total of 318,063 cells across six tissue sections (Figures S1-S3). For the Xenium DG dataset, this resulted in 25,387 cells discarded from the total of 532,331 cells across four tissue sections (Figures S8-S10).

Feature selection typically follows after normalization in scRNA-seq analyses to facilitate computation due to the large number of genes. However, the Xenium datasets only contain a few hundred to a few thousand genes, allowing this step to be bypassed and all gene features to be used in downstream analyses. Features representing negative controls were excluded from downstream analyses as they do not provide additional biological information.

#### 4.2.3 Spatial domain annotation of Xenium dlPFC data

The human Visium dlPFC dataset [33] was used as the reference for label transfer with spaTransfer to the Xenium dlPFC data as the target. A rank of 25 was used for the number of NMF factors. Spatially-aware label smoothing was applied using a 10 nearest neighbors graph.

#### 4.2.4 Spatial domain annotation of Xenium DG data

The human Visium DG dataset [39] was used as the reference for label transfer with spaTransfer to the Xenium DG data as the target. A rank of 100 was specified for the number of NMF factors choosen. Spatially-aware label smoothing was then run on the transferred labels in order to obtain more contiguous domains and reduce noise in the Xenium annotations. For each sample, a 15 nearest neighbors graph was constructed based on spatial coordinates and label smoothing was performed across this graph (see Methods).

### 4.3 Overview of spaTransfer

#### 4.3.1 Motivation for using non-negative matrix factorization

The primary motivation behind using NMF in spaTransfer was its ability to decompose a dataset into its constituent parts, which are interpretable as standalone factors, allowing for the identification of signal that is conserved between different experimental platforms. For example, a group may have previously generated Visium data for their tissue of interest and now want to use their Visium dataset to annotate a newly generated Xenium dataset. The observational unit in the Visium platform known as a “spot” is a circular region spanning 50 microns in diameter. Due to its size, each spot typically contains multiple cells and users are only able to observe the transcriptional profile of the entire spot rather than the individual cells within. In contrast, Xenium is an imaging-based platform that offers sub-cellular resolution measurements of gene expression, and observations can be aggregated to the single cell level. To annotate a single-cell resolution dataset with the spatial domain labels from a spot resolution dataset, spaTransfer identifies latent factors in the reference atlas that represent biological processes of interest, in this case spatial domains. The same set of latent factors are then computed in the unlabeled target dataset, but at a single-cell resolution. Previous work has shown that this transfer of latent factors across datasets of different resolutions accurately preserves spatially organized biological signals [27, 28]. Similar to previous work, we found here that spaTransfer is able to leverage the preserved information encoded by the latent factors to transfer labels between datasets of different features and resolutions.

#### 4.3.2 Using NMF to identify latent factors in reference atlas

NMF was originally proposed by Lee and Seung [61] as an alternative to PCA and vector quantization. The original paper found that NMF applied to images of faces resulted in a decomposition into factors representing individual facial landmarks such as the nose, mouth, or eyes. Similar to PCA, NMF factorizes a matrix into two lower-rank non-negative matrices *W* and *H*, but with the additional constraint of *W* and *H* being matrices with non-negative entries. Additionally, the factorization can instead solve for *W* and *H* in the following equation:

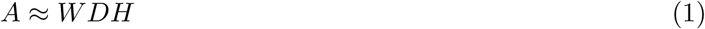

Where *D* is a scaling factor that ensures each NMF factor sums to 1. spaTransfer uses the *singlet* R package [62] which relies on the NMF implementation in RcppML [23]. If *A* is a gene expression matrix with log-normalized gene expression counts from single-cell or SRT experiments with each row representing a gene and each column representing a cell, then *W* is a gene by rank matrix and *H*is a rank by cell matrix. Similar to PCA, W can be thought of as a loadings matrix and *H* is a factors matrix, with each row in *H* representing a factor. The non-negativity constraint on the factorization yields factors that can be interpreted additively and independently of other factors, unlike in PCA [61]. Thus, when an appropriate rank is used for the factorization, NMF factors can be interpreted as representing a particular spatial domain or cell type.

Briefly, the NMF model is fit using alternating least squares. First, *W* is randomly initialized and a least squares procedure is used to find an estimate for *H*. Then, least squares is used again with *H* held constant to find a better estimate for *W*. This process is repeated until the relative change of and between iterations is smaller than the user-specified tolerance parameter.

When performing NMF on a reference dataset, spaTransfer offers two options for specifying the rank of the factorization *L*. The first option is for the user to provide *L* based on domain knowledge of the tissue. For the purposes of label transfer, we recommend that at minimum *L* should be equal to the number of levels in the annotation. This provides a necessary, but not sufficient condition for each level of the annotation to be uniquely represented by a NMF factor. However, it has been previously observed that NMF factors can also represent technical artifacts [23, 26–28] and thus a higher value of *L* may be needed to ensure representation of all annotation levels. Conversely, a value of *L* that is too high can result in overfitting and poor prediction performance. The second option presented by spaTransfer aims to address this issue. If the user does not specify a value for *L*, then by default spaTransfer will run automatic rank determination using the RunNMF() function from the singlet package [62]. The automatic rank determination approach uses cross validation to select an optimal value for *L* while accounting for overfitting. Specifically, the automatic rank determination function will “mask” a user-specified percentage of the entries in the matrix *A*. By default, this is set to 5% of the total number of entries in *A*. These values are masked by replacing them with random normal variables. An NMF model is then fit at low value of *L*. The performance of the NMF model at this low rank is then evaluated using the mean squared error between the entries that were previously masked and the model’s predicted values for those entries. This process is repeated for different values of until a value that minimizes the mean squared error is found. By default, the automatic rank determination function runs three independent replicates of this procedure to assess consistency of results across different initializations.

#### 4.3.3 Transfer of NMF factors to target datasets

Once factors are extracted from the reference dataset (*A*), the gene by rank loadings matrix (*W*) is used to predict NMF factors on the expression values of the target dataset (*A*^′^). The prediction can be expressed as solving for *H*^′^ in the following equation:

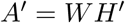

where *A*^′^ is the log-normalized counts matrix of the target dataset and *W* is the loadings matrix fitted on the reference dataset. Then, the least-squares estimate of *H*^′^ can be computed as follows by solving the following equation

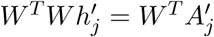

for each *h*^′^*_j_*, where *h_j_*^′^ refers to the *j*th column of *H*^′^. This was done using the project function from RcppML [23] following previously established workflows [27, 28].

Writing the reference factorization as

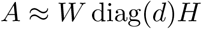

singlet normalises each row of *H* to sum to one, so that for every factor *i*

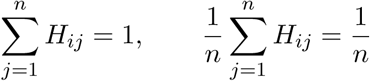

where *n* is the number of reference spots. We then fit the multinomial model on *H^T^* which therefore learns thresholds for classes in the units of *H^T^*. However, project does not normalize *H*^′^, so values in *H*^′^*^T^* are not guaranteed to be in the same range as that of *H^T^*, hindering the predictive ability of the multinomial model. To normalize *H*^′^*^T^* to the same units as *H^T^*, we define *δ* ∈ R*^k^* and use it to scale *H*^′^ before feeding the projected factors into the multinomial model:

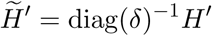

Let *ρ* denote the fraction of reference genes also measured in the target. When *ρ* ≥ 0.5 (or at least 50% of the genes are measured in the target), then we take *δ* = *d*. When *ρ <* 0.5 (or fewer than 50% of the genes are measured), then the reference *d* no longer describes the target, and *δ* is estimated from the target itself. Letting *m* be the number of target cells,

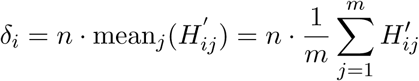

This can be rewritten so that the mean weight per target cell equals the reference mean weight per spot:

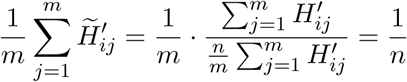

Intuitively, it expresses each cell’s weight as a fraction of the total target weight for that factor. Then, it multiplies by *m/n*. This ensures

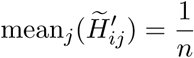

matching the reference factor’s mean. This transformation matches the mean weight of each target factor to its reference mean of 1*/n*, placing the projected factors on a comparable scale for prediction with the fitted multinomial models.

In the case of cross-platform label transfer, the number of genes often differs between the reference and the target datasets, making matrix multiplication impossible due to incompatible matrix dimensions. To address this issue, the loadings matrix is subsetted after fitting NMF to contain only the genes that are found in both the reference and target datasets. The subsetted loadings matrix is then used in the project function from RcppML [23] to predict factors in the target dataset. In the case of the Xenium DG label transfer with the Visium DG dataset as the reference, we subset the original full-transcriptome loadings matrix to only the 5000 genes that are in the Xenium data. To ensure that the predicted factors are still meaningful, we simulated fitting NMF on randomly generated matrices and reconstructing factors using a subsetted loadings matrix. These simulations showed that subsetting to 10% of the original features will preserve the distribution of the majority of the factors, and subsetting to 25% of the original features preserves the distribution of all factors (Figure S18), assuming that all features are equally informative. Thus, retaining 5000 out of 20,000 genes should be sufficient for predicting factors in the target dataset.

#### 4.3.4 Multinomial annotation prediction model

spaTransfer uses a multinomial model to generate annotation predictions on the target dataset with the annotation as the response as the selected NMF factors as the covariates. The multinomial model is implemented through the *glmnet* R package [25]. Let *C* denote the cell type or spatial domain annotations in the reference dataset. For a given observation in class *c* in *C*, we model

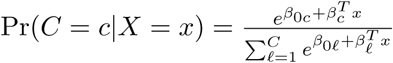

where *x* denotes the NMF factors for the observation. This yields a linear predictor fit for each class of the annotations. Elastic net regularization [24] implemented in the *glmnet* R package [25] is used to prevent overfitting, resulting in the following log-likelihood:

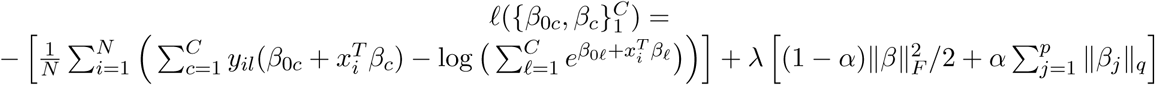

Where *y_il_* is an indicator denoting whether observation *i* is a member of class *l*, *α* is the elastic net parameter, controlling the weighting on the lasso and ridge penalties, and *λ* controls the strength of the penalty term. The parameter *q* controls the type of lasso penalty used. If *q* is set to 1, each of the *K* × *p* coefficients each have their own lasso penalty, and can be zero or non-zero independent of other coefficients. If *q* is set to 2, the grouped lasso penalty is used and the *K* coefficients corresponding to the same variable will all be either zero or non-zero. We chose to use the grouped lasso penalty to ensure the same set of factors were selected across each of the classes. Further details of the model-fitting procedure can be found in glmnet [25].

#### 4.3.5 Label smoothing

After annotations have been transferred into the target datasets, spaTransfer provides an optional spatially-aware smoothing algorithm to increase coherency of transferred domains. The foundation of the smoothing algorithm follows a previously established method [31] which builds a *k*-nearest neighbors (kNN) graph based on the spatial coordinates for each cell using the *dbscan* R package [63]. The number of neighbors is specified by the user. Then, for each cell, the proportion of neighbors labeled with each level in the annotations is computed. If any of the level proportions in the neighborhood is greater than the user-specified threshold, the cell is then assigned to the level with the greatest proportion. Otherwise, the cell retains its original label. This process is repeated for a user-specified number of iterations, with the number of cells being reassigned generally decreasing with each iteration. Our modification to this algorithm allows for a different threshold to be used for each level in the annotations, rather than having the same threshold for all levels. This is motivated by the different cell densities exhibited by spatial domains.

### 4.4 Downstream analyses of spatial domains

#### 4.4.1 Pseudobulk processing of DG data

After annotating each cell’s spatial domain identity using spaTransfer, we sought to explore the differences between spatial domains across the lifespan. To this end, we summed up the raw counts for each gene across all cells of each spatial domain using the aggregateAcrossCells() function from the *scuttle* R package [60]. This resulted in 40 pseudobulked samples (4 donors x 10 spatial domains) with each one representing a unique donor-spatial domain combination. The choroid plexus (CP) pseudobulk samples were removed as the CP is a secretory tissue that has been shown to account for the majority of variation when included in PCA [39]. The pseudobulk samples representing the stratum radiatum (SR) domain were also removed as we found that they also drove PC1 (57% of variance explained) (Figure S14). We found a similar result with the pseudobulk samples representing the CA1 and SL regions in the teen donor. This is likely due to the CA1 region being absent in the tissue section and thus any cells with the CA1 label are misclassified. Because only a small number of cells had these labels (197 CA1 cells, 168 SL cells), the teen CA1 and SL pseudobulk samples were also excluded from downstream analyses. This procedure was done separately for each donor and the pseudobulked samples were combined into one large SpatialExperiment [59] object, with each pseudobulk sample (column) representing a unique donor-domain combination. Genes that are lowly expressed in the spatial domains are removed from the pseudobulked samples using filterByExpr() in the *edgeR* R package [64]. Finally, log-normalized counts were calculated from the pseudobulk samples using calcNormFactors() in *edgeR*. PCA was used to identify the primary sources of variation in the pseudobulked dataset using the log-normalized counts computed above using runPCA from *scater* [60] using 10 PCs.

### 4.5 Comparison against existing methods

#### 4.5.1 Comparing accuracy of predicted labels

To evaluate the performance of spaTransfer, we ran three existing single-cell reference mapping algorithms. Specifically, these were singleR [14], Seurat (v5) [10, 50], and scmap [15]. We benchmarked the performance of these methods on the Visium dlPFC dataset [33]. Half the samples were used as the reference atlas and the remaining half were used as the target samples. Each of the methods were then used to predict spatial domain annotations on the target samples. To quantify the performance of each method, the adjusted Rand index (ARI) [65, 66] and normalized mutual information (NMI) [66] were computed between the labels annotated by the original authors of the manuscript in the reference and the predicted labels from spaTransfer or the other three algorithms (singleR, Seurat, and scmap).

#### 4.5.2 Comparing spatial contiguity and internal validity of predicted annotations

Due to the lack of ground truth labels for the Xenium datasets, we were unable to compute external evaluation metrics to evaluate the accuracy of transferred labels. Thus, we compared the performance of spaTransfer to BANKSY [31], MENDER [51], PRECAST [52], and SLAT [53] by computing metrics quantifying the spatial contiguity and internal validity metrics of transferred spatial domain labels (described below). All of these methods perform unsupervised and reference-free spatial domain detection, with the exception of SLAT. SLAT is a reference mapping method that uses a Visium dataset as the reference, and transfers labels from the Visium dataset to a target Xenium dataset. However, since it is only designed for the Visium to Xenium setting, we chose to benchmark it along with the other unsupervised methods due to the lack of Xenium ground truth labels.

For each of the methods, we computed the proportion of abnormal spots (PAS) [55] and the CHAOS metric [55] to evaluate the spatial contiguity of the predicted annotations. The PAS is defined as the number of abnormal cells out of the total number of cells in a tissue section. An abnormal cell is defined as a cell that has a different spatial domain label that is different from 6 or more of its nearest 10 neighbors [55]. The CHAOS metric is computed for each tissue section by first creating a 1-nearest neighbor graph for each spatial domain using the spatial locations of each cell. Then, for each pair of cells in domain *r*,

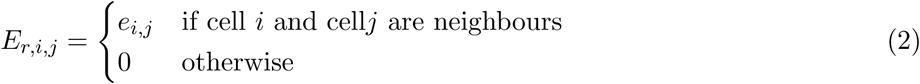

and

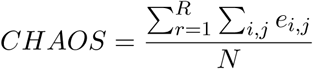

where *R* is the total number of spatial domains and *N* is the total number of cells in the tissue section [55]. Finally, the Silhouette index was used to evaluate the internal validity of clusters. For each tissue section, PCA was first computed on the log-normalized counts using the *scanpy* Python package [67] with default parameters. The Silhouette score was then computed using the *scikit-learn* package [66], with euclidean distance as the metric.

#### 4.5.3 Inputs and settings for the methods

We applied the two existing label transfer methods to our Xenium dlPFC dataset. For singleR and Seurat, we supplied the Visium dlPFC dataset as the reference. For singleR and Seurat, we followed the author-recommended preprocessing workflows provided in each method’s tutorials. For spaTransfer, we used the log2-normalized gene expression and a factorization rank of 25. Batch correction was not performed prior to running any of the methods. We used the same set of reference and query datasets for all three methods, thus allowing for a fair comparison between methods. For all three methods, we performed quality control and pre-processing of our datasets as described above and then followed the author-provided tutorials and their recommended parameter settings.

We also applied four domain detection methods that are applicable to imaging based SRT to our data. For BANKSY [31], MENDER [51], PRECAST [52], and SLAT [53], we provided the Xenium dlPFC dataset as the input for each method, and followed the authors’ tutorials to preprocess the data and select parameters.

### 4.6 Rank and factorization algorithm sensitivity analysis

To assess the sensitivity to the choice of rank for NMF, we ran spaTransfer on ranks of 5, 10, 15, 20, 25, 40, 60, 100, 150, 200, 300, 400, and 500. The data used for this assessment was the same Visium dlPFC dataset described above. The same reference-target split was used as the benchmark analyses. Using the Visium BayesSpace labels as the ground truth, we quantified the accuracy of predictions at each rank using the ARI. To quanitfy the accuracy of the matrix reconstruction, we used the cross_validate_nmf function from *singlet* which reported the cross-validation mean squared error on held out entries.

To assess performance differences between NMF and PCA for the spaTransfer prediction workflow, we compared the two using the same workflow for the dlPFC Visium to Xenium label transfer described above. For spaTransfer, we ran the same workflow as above, namely performing the factorization at a rank of 25. For PCA, we preserved all parts of the spaTransfer method except we replaced the NMF step with PCA at a rank of 25 to match the NMF workflow. We used absolute Pearson correlation between the factors from the two method and the Visium ground-truth BayesSpace labels.

### 4.7 Software and data availability

spaTransfer is freely available as an R package on GitHub (github.com/cindyfang70/spaTransfer). Code to reproduce all preprocessing, analyses, and figures in this manuscript is available from GitHub at github.com/LieberInstitute/spatransfer-manuscript. We used spaTransfer version 0.0.0.9000 for the analyses in this manuscript. Analyses were performed with R version 4.3. Data visualizations were performed with *ggplot2* [68] and *escheR* [69] R packages. The processed data associated with the analyses in this manuscript are available at zenodo.org/records/17872293. The raw Xenium data files are currently being uploaded to GEO and the link will be shared at github.com/LieberInstitute/spatransfer-manuscript.

## Acknowledgments

Portions of some figures were created with BioRender.com. We thank 10x Genomics for reagents and early access to a precommercial version of the 5000-Plex Xenium Prime assay. The authors extend their gratitude to donor families who contributed tissue and clinical information for this study. We would also like to thank Amy Deep-Soboslay and James Tooke at the Lieber Institute for Brain Development for their exceptional efforts in the curation of the brain samples for this investigation. We thank the maintainers of the Joint High Performance Computing Exchange (JHPCE) compute cluster at Johns Hopkins Bloomberg School of Public Health for providing essential computing resources. We also thank Hicks lab for their constructive comments and suggestions. We thank Yi Wang for suggesting the idea of a multinomial model to predict labels.

## Author Contributions Statement

Xenium data was generated by ADR, KDM, SM, supervised by SCP, KRM, KM. RM preproccessed the Xenium data through SpaceRanger. CF conceptualized the spaTransfer methodological framework with suggestions from SCH. The formal analysis and benchmarking was led by CF with suggestions from BG. Figures were designed by CF. Software was developed by CF with input from other co-authors. SCP, KMaynard KMartinowich, and SCH administered and supervised the project. CF and SCH wrote the original draft of the text and all co-authors edited and approved the final manuscript.

## Supplementary Tables

**Supplementary Table S1:**
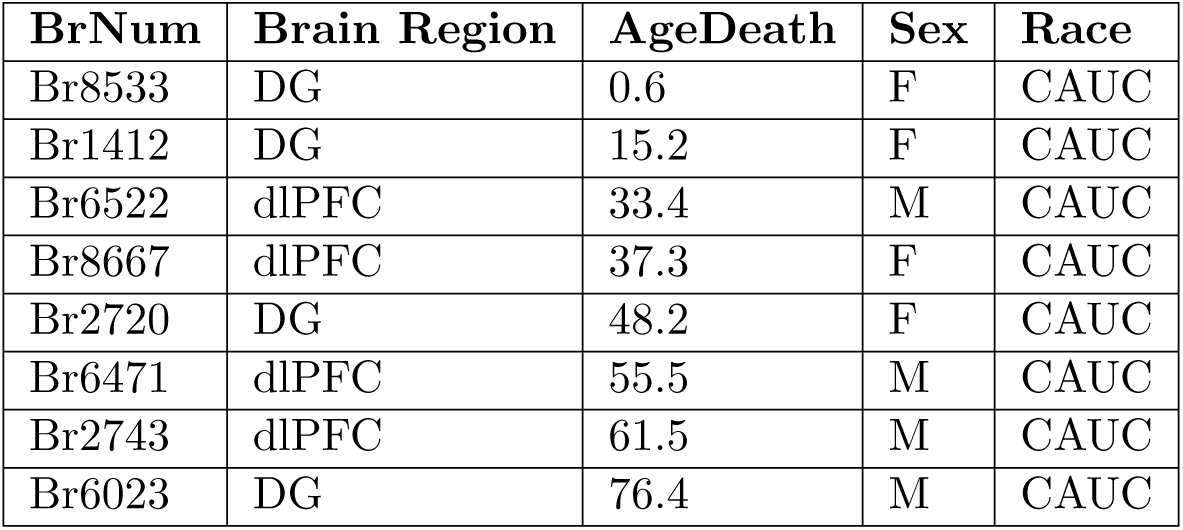
Demographic information on brain donors. Demographic information for all brain donors in this study, including donor ID (BrNum), brain region, age at time of death, sex, ancestry, screening RIN, performed in prefrontal cortex (PFC) at time of tissue collection, postmortem interval (PMI) and psychiatric diagnosis.

## Supplementary Figures

**Supplementary Figure S1:**
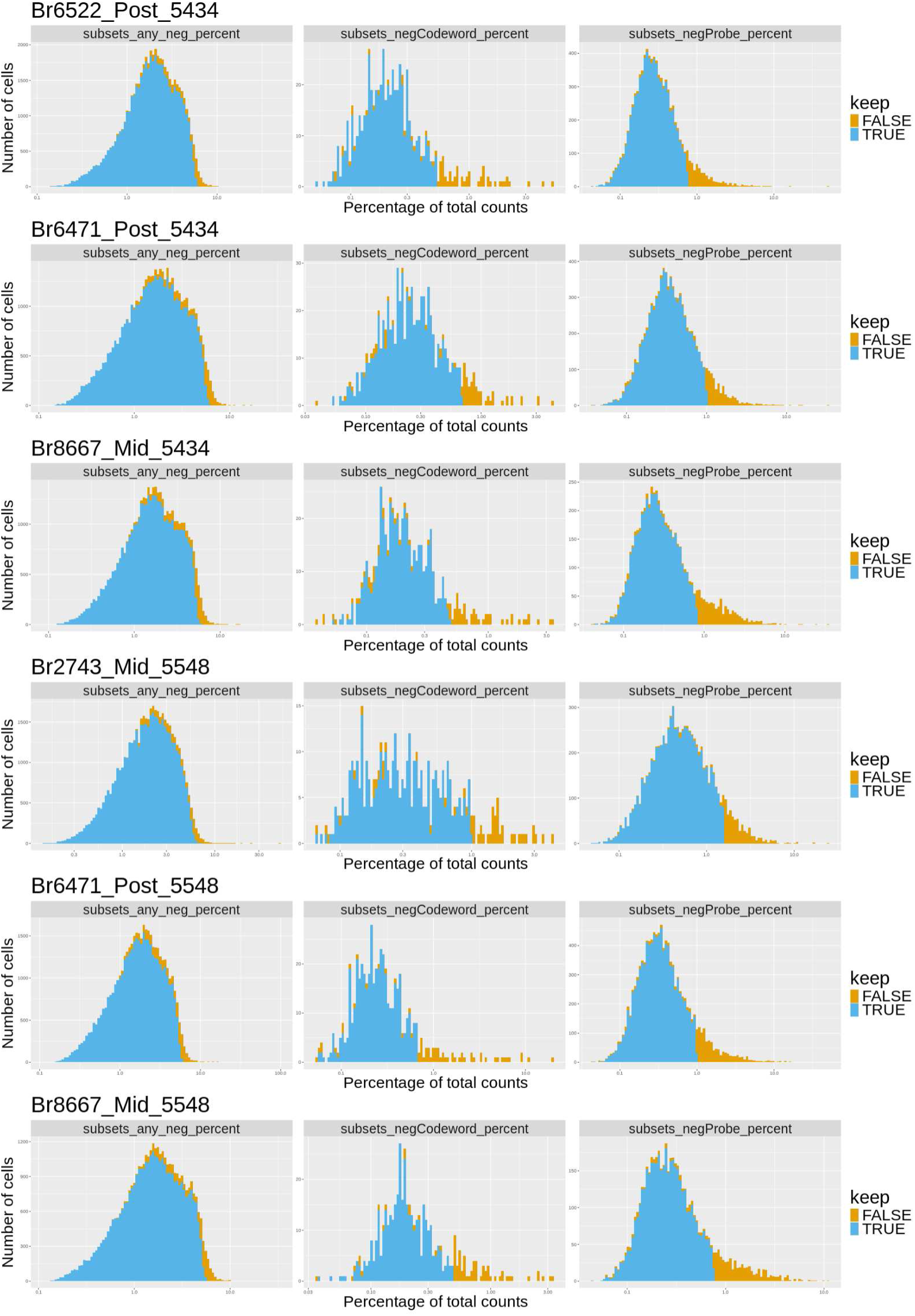
Distribution of quality control metrics used to filter out low-quality spots for dlPFC samples. Three quality control (QC) metrics were considered: (i) the percent of reads mapping to any negative control codewords or probes (column 1), (ii) the percent of reads mapping to any negative control codewords (column 2), and (iii) the percent of reads mapping to any negative control probes (column 3). Rows correspond to the six Xenium tissue sections. Distributions of the QC metrics are provided for the cells that were kept (blue) or that were removed (yellow).

**Supplementary Figure S2:**
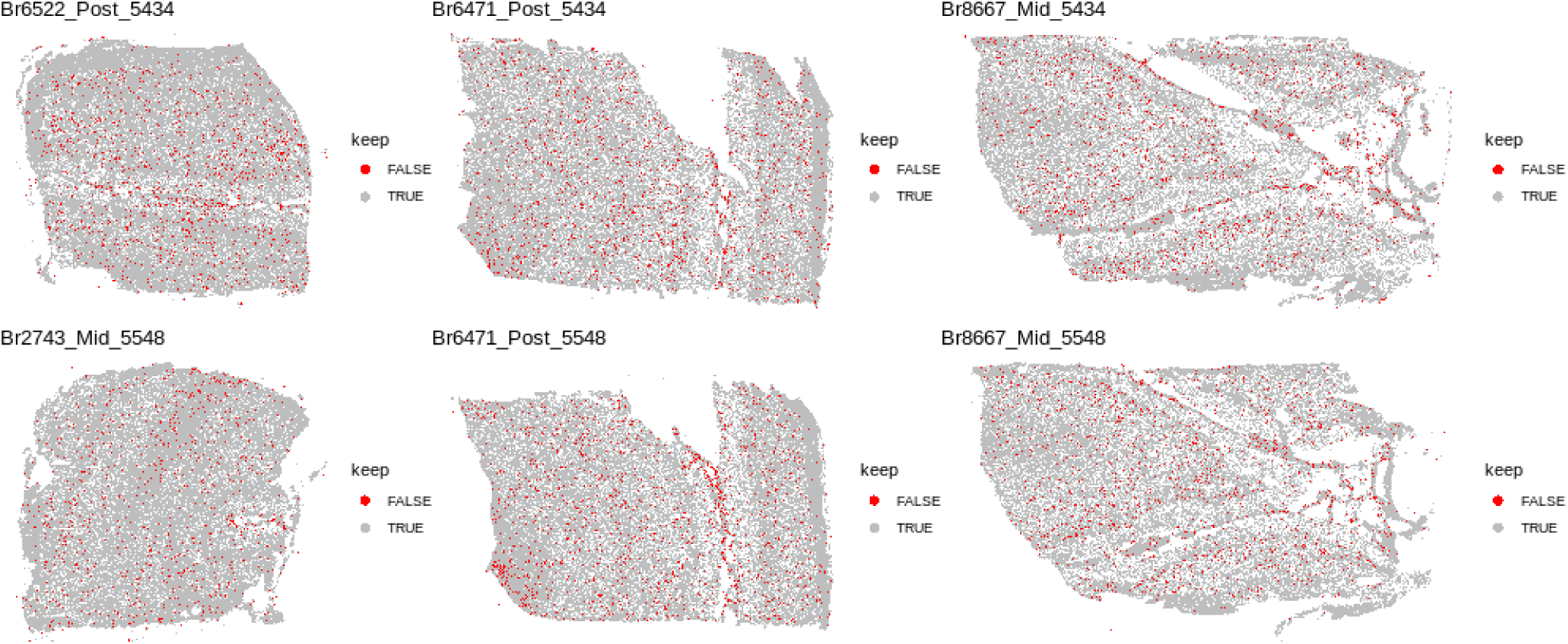
Spotplots of discarded cells based on QC metrics. Six cell-level tissue plots of discarded cells (red) for each Xenium dlPFC capture area following quality control. Cells were discarded based on the percentages of reads from negative control probes and negative control codewords.

**Supplementary Figure S3:**
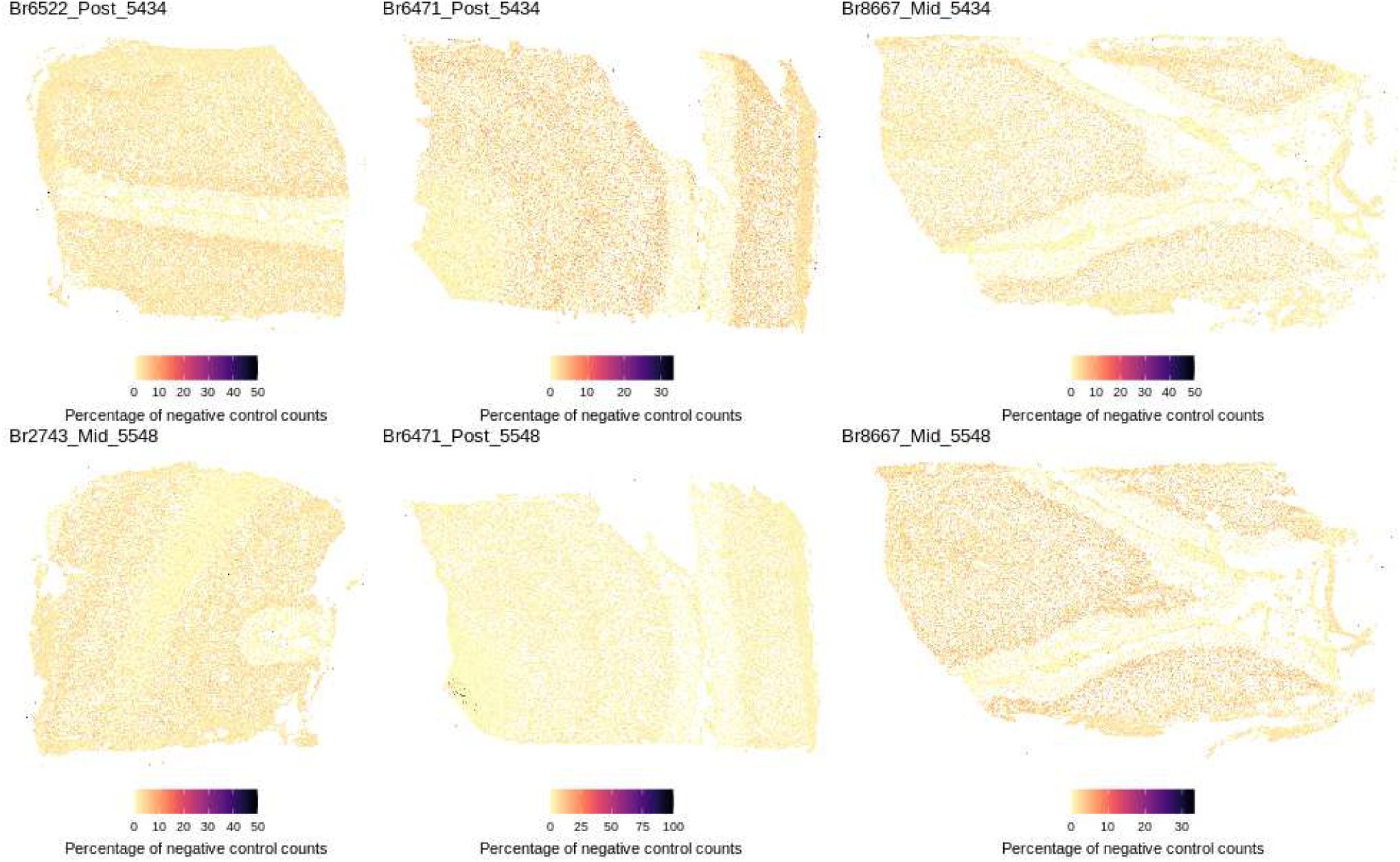
Spotplots of expression counts of negative control features. Six cell-level tissue plots of the expression of negative control features per cell for each Xenium dlPFC capture area. Counts from both negative control probes and negative control codewords were aggregated into one feature and plotted on the tissue.

**Supplementary Figure S4:**
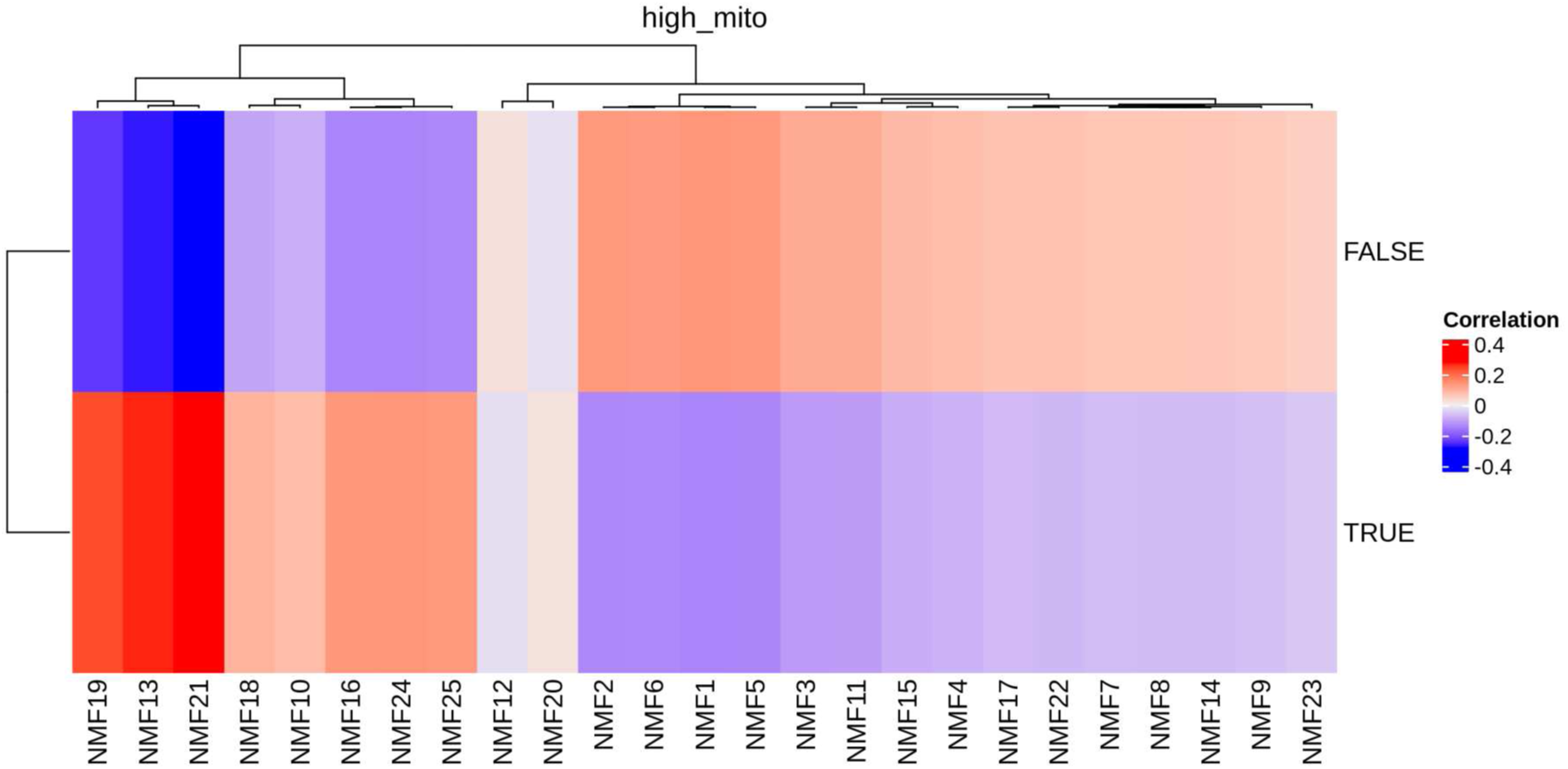
Heatmap showing Pearson correlation between NMF factors and spots labeled as having high or low percentages of mitochondrial counts. NMF19, NMF13, and NMF21 are highly correlated with spots with high mitochondrial percentages.

**Supplementary Figure S5:**
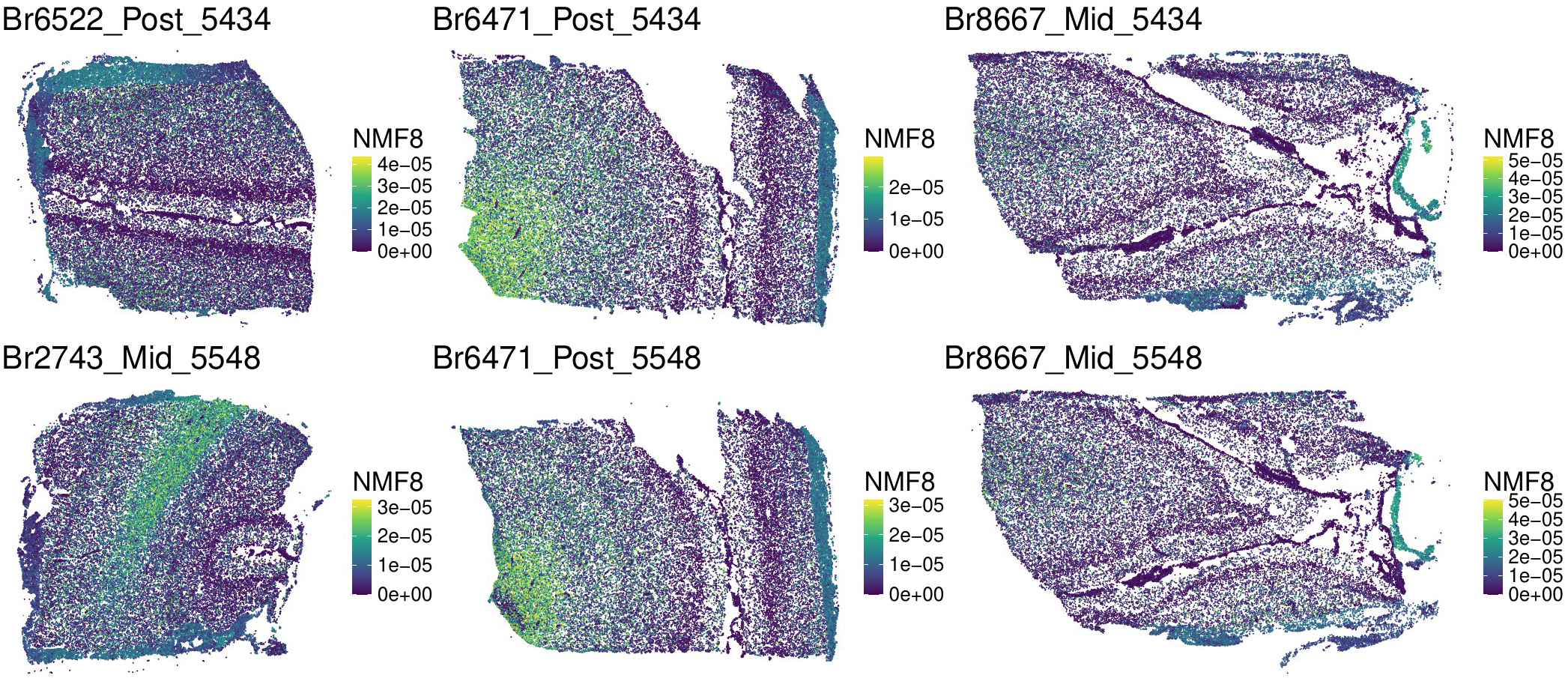
Spotplots of NMF8 weights (learned from the loading matrix from the reference dataset) visualized in the target dataset. Six cell-level tissue plots of the NMF8 weights (learned from the loading matrix from the 10x Genomics Visium reference dataset) visualized for each Xenium dIPFC capture area.

**Supplementary Figure S6:**
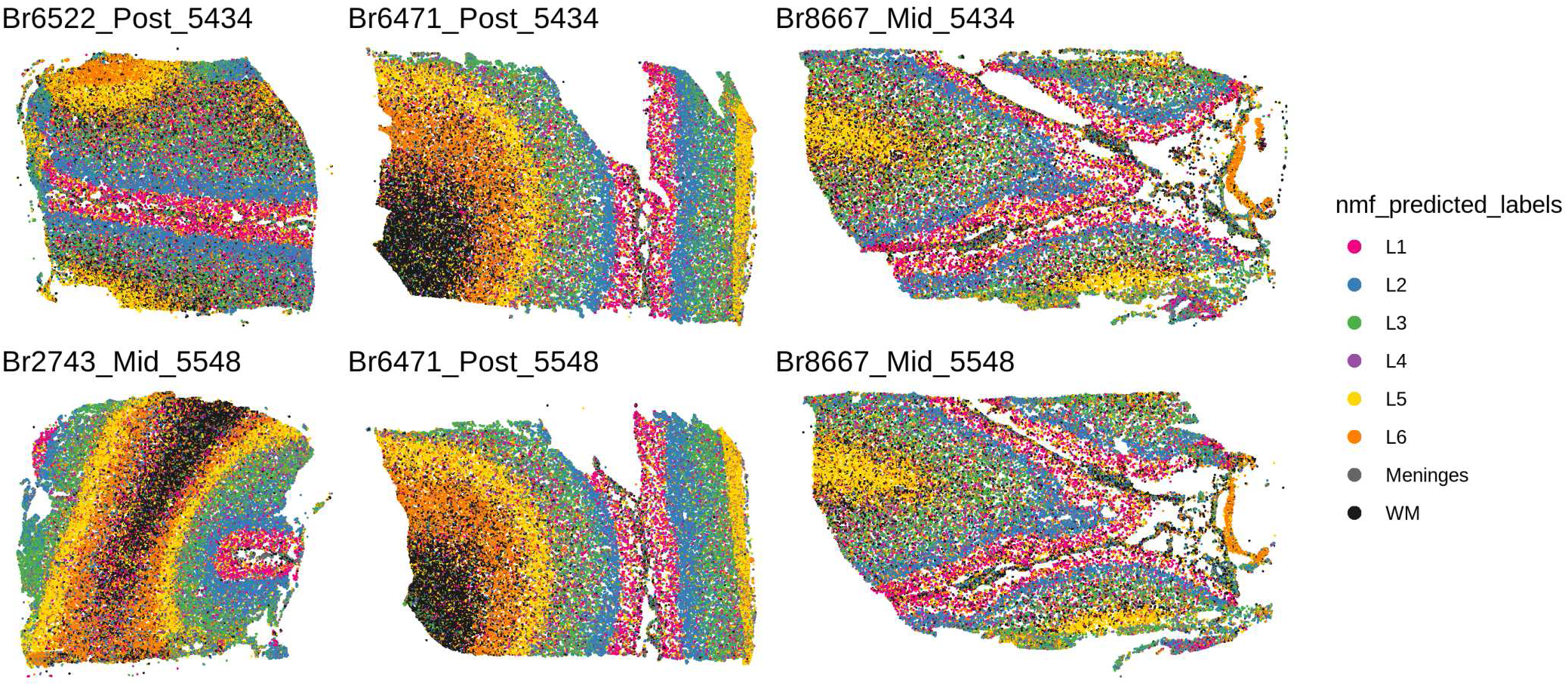
Spotplots of predicted spatial domain labels from spaTransfer (unsmoothed). For each of the six Xenium dlPFC capture areas, the cells are colored by predicted layer identity. No smoothing has been applied and these are the raw predicted labels.

**Supplementary Figure S7:**
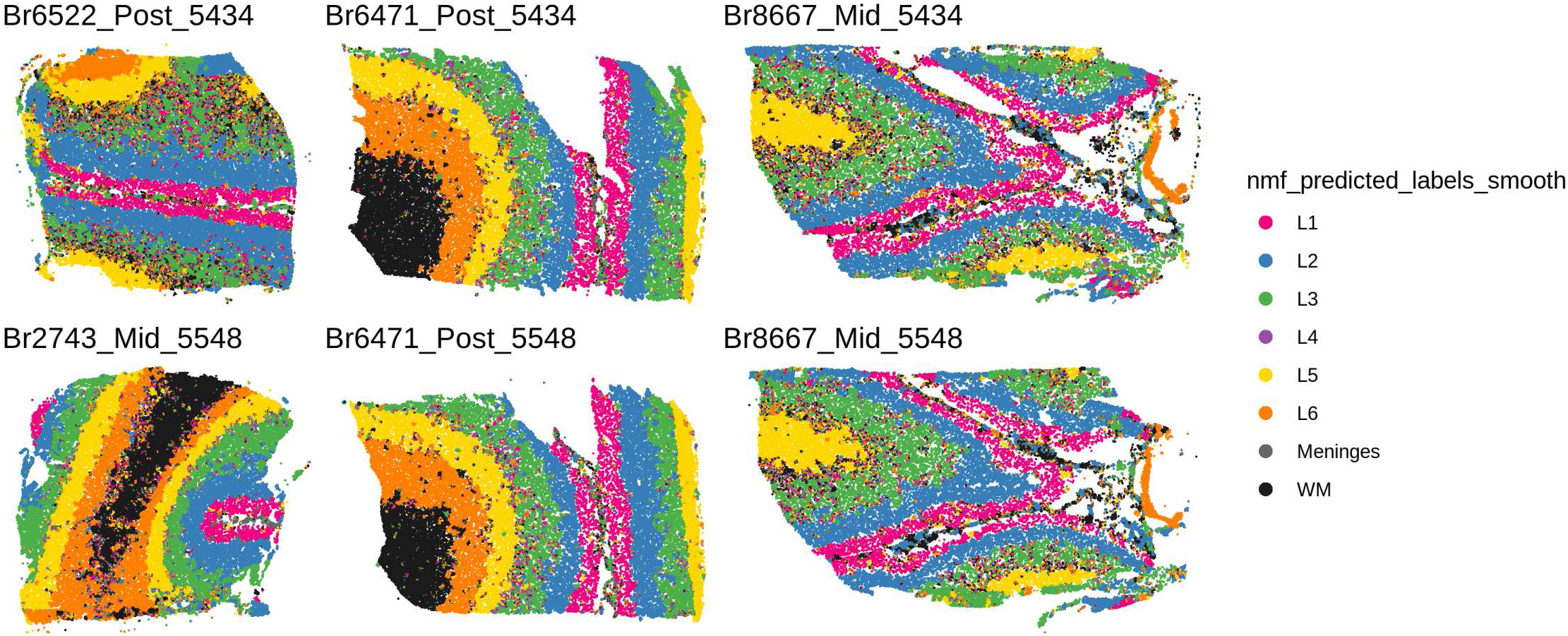
Spotplots of predicted spatial domain labels from spaTransfer (smoothed). Similar to Figure S6, but smoothing (see Methods) has been applied to the raw predicted labels.

**Supplementary Figure S8:**
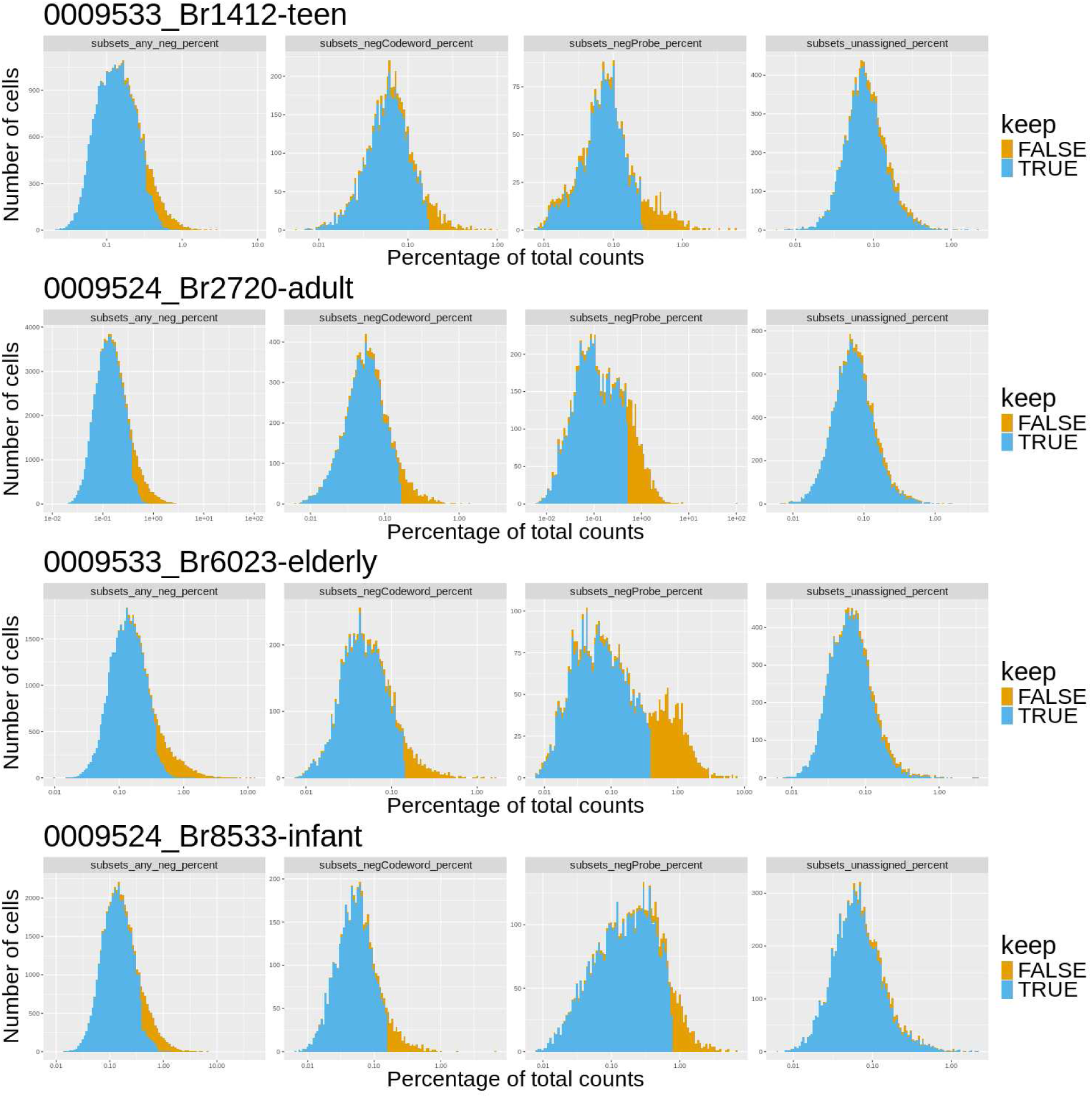
Distribution of quality control metrics used to filter out low-quality spots for HPC samples. Four quality control (QC) metrics were considered: (i) the percent of reads mapping to any negative control codewords or probes (column 1), (ii) the percent of reads mapping to any negative control codewords (column 2), (iii) the percent of reads mapping to any negative control probes (column 3), and (iv) the percent of reads mapping to any unassigned codewords. Rows correspond to the four Xenium tissue sections. Distributions of the QC metrics are provided for the cells that were kept (blue) or that were removed (yellow).

**Supplementary Figure S9:**
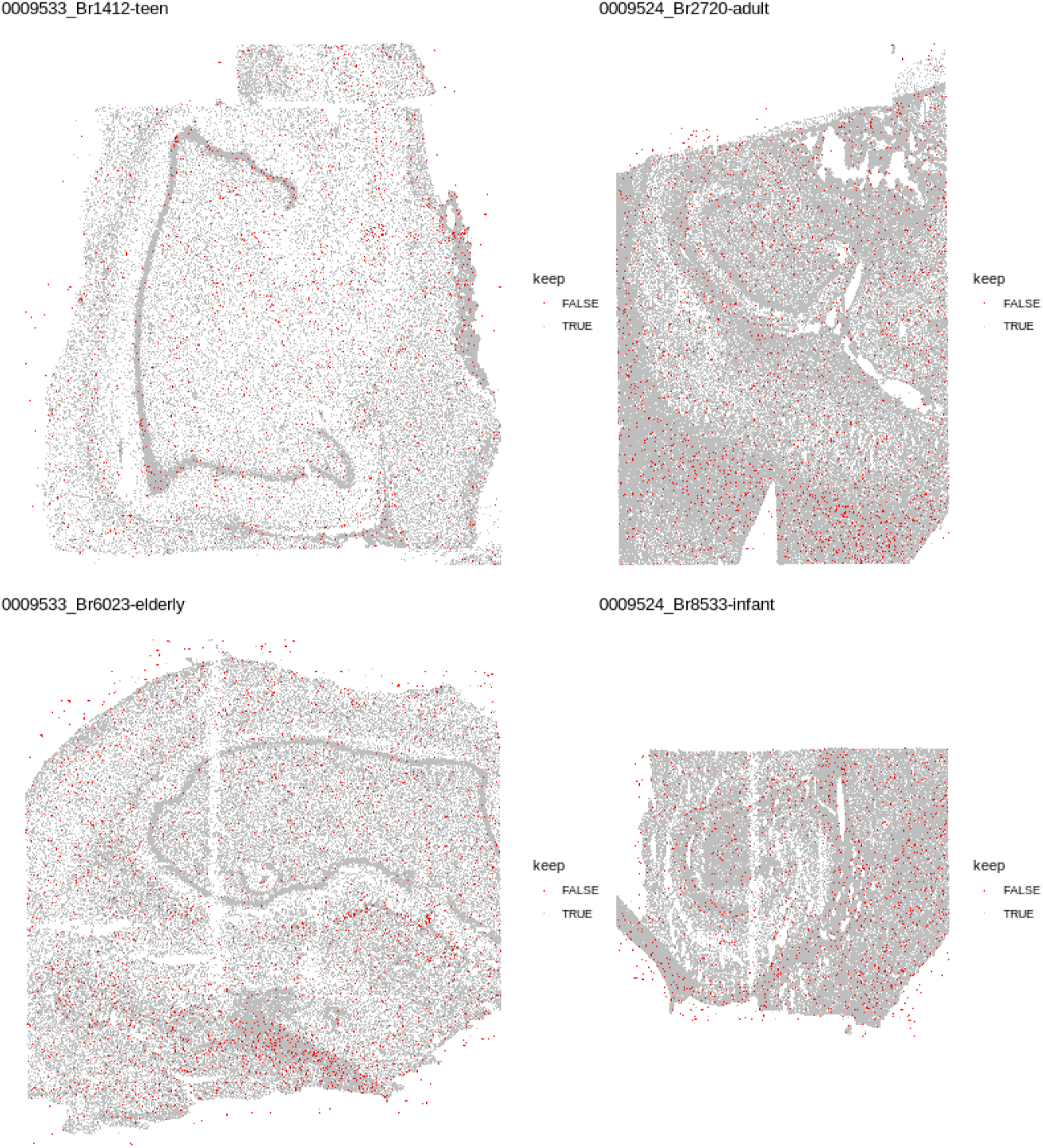
Spotplots of discarded cells based on QC metrics. Four cell-level tissue plots of discarded cells (red) for each Xenium HPC capture area following quality control. Cells were discarded based on the percentage of reads from negative control features.

**Supplementary Figure S10:**
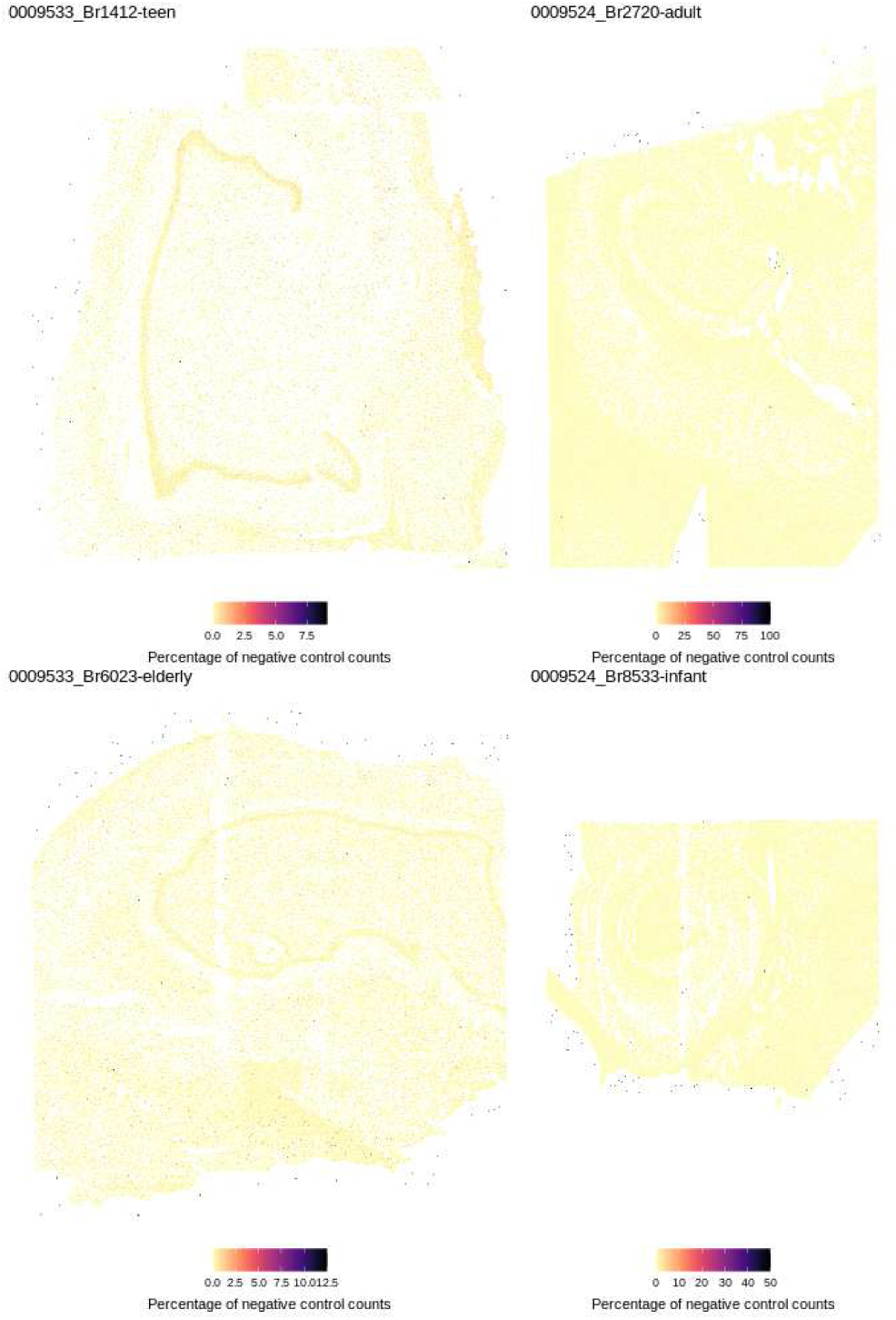
Spotplots of expression counts of negative control features. Four cell-level tissue plots of the expression of negative control features per cell for each Xenium HPC capture area. Negative control probes, negative control codewords, and unassigned codewords were aggregated into one feature and plotted on the tissue.

**Supplementary Figure S11:**
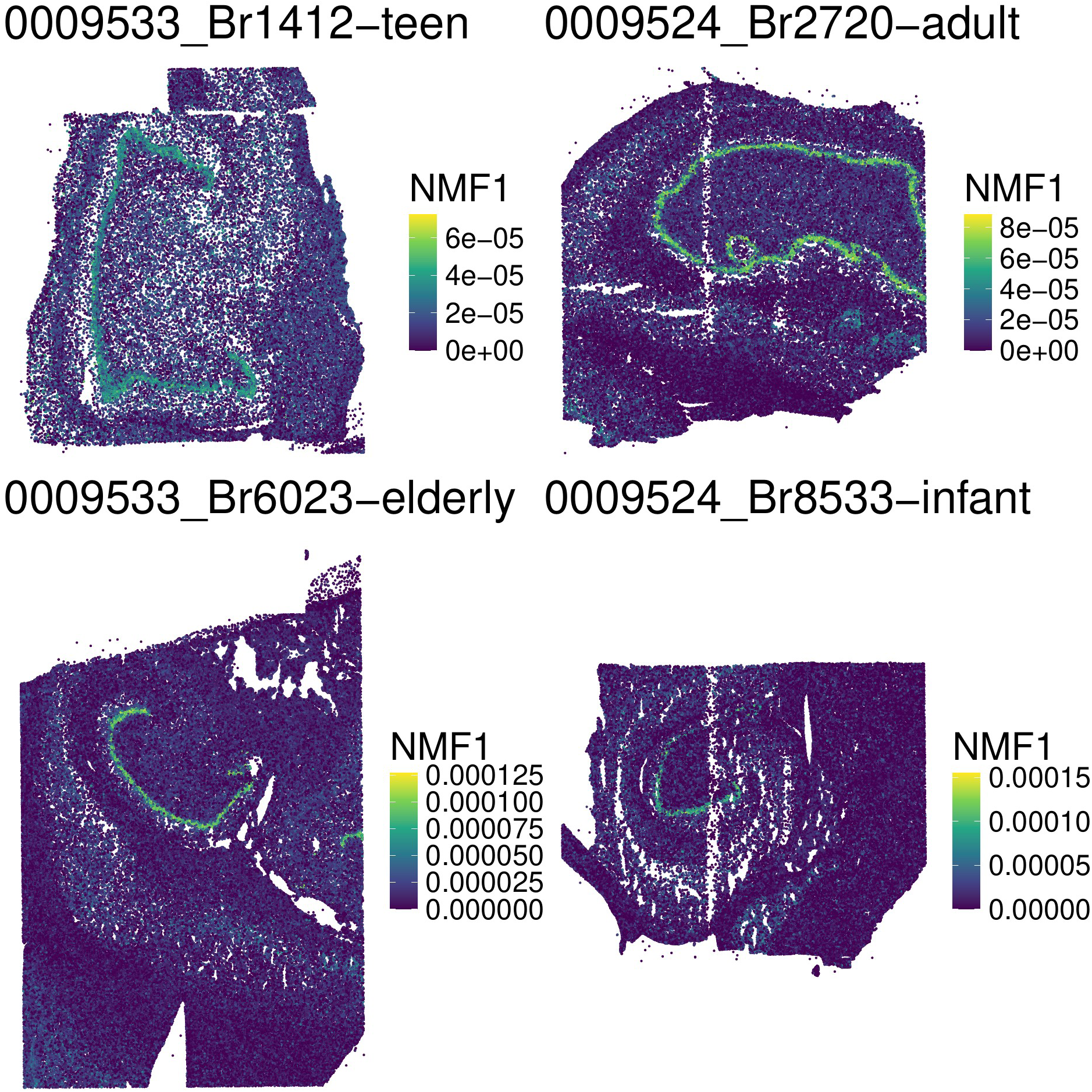
Spotplots of NMF1 weights (learned from the loading matrix from the reference dataset) visualized in the target dataset. Four cell-level tissue plots of the NMF1 weights (learned from the loading matrix from the 10x Genomics Visium reference dataset) visualized for each Xenium HPC capture area.

**Supplementary Figure S12:**
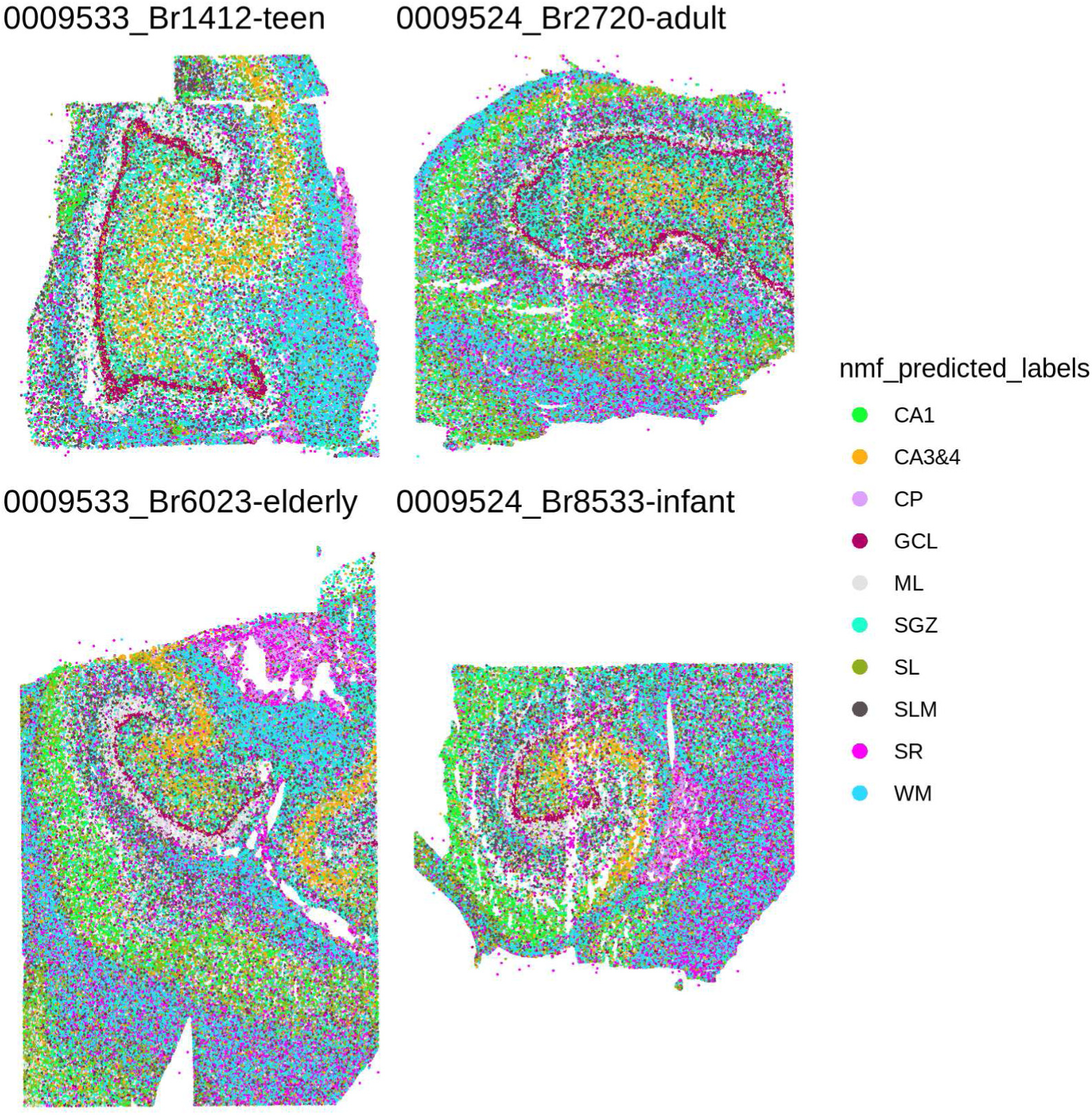
Spotplots of predicted spatial domain labels from spaTransfer (unsmoothed). For each of the four Xenium HPC capture areas, the cells are colored by predicted layer identity. No smoothing has been applied and these are the raw predicted labels.

**Supplementary Figure S13:**
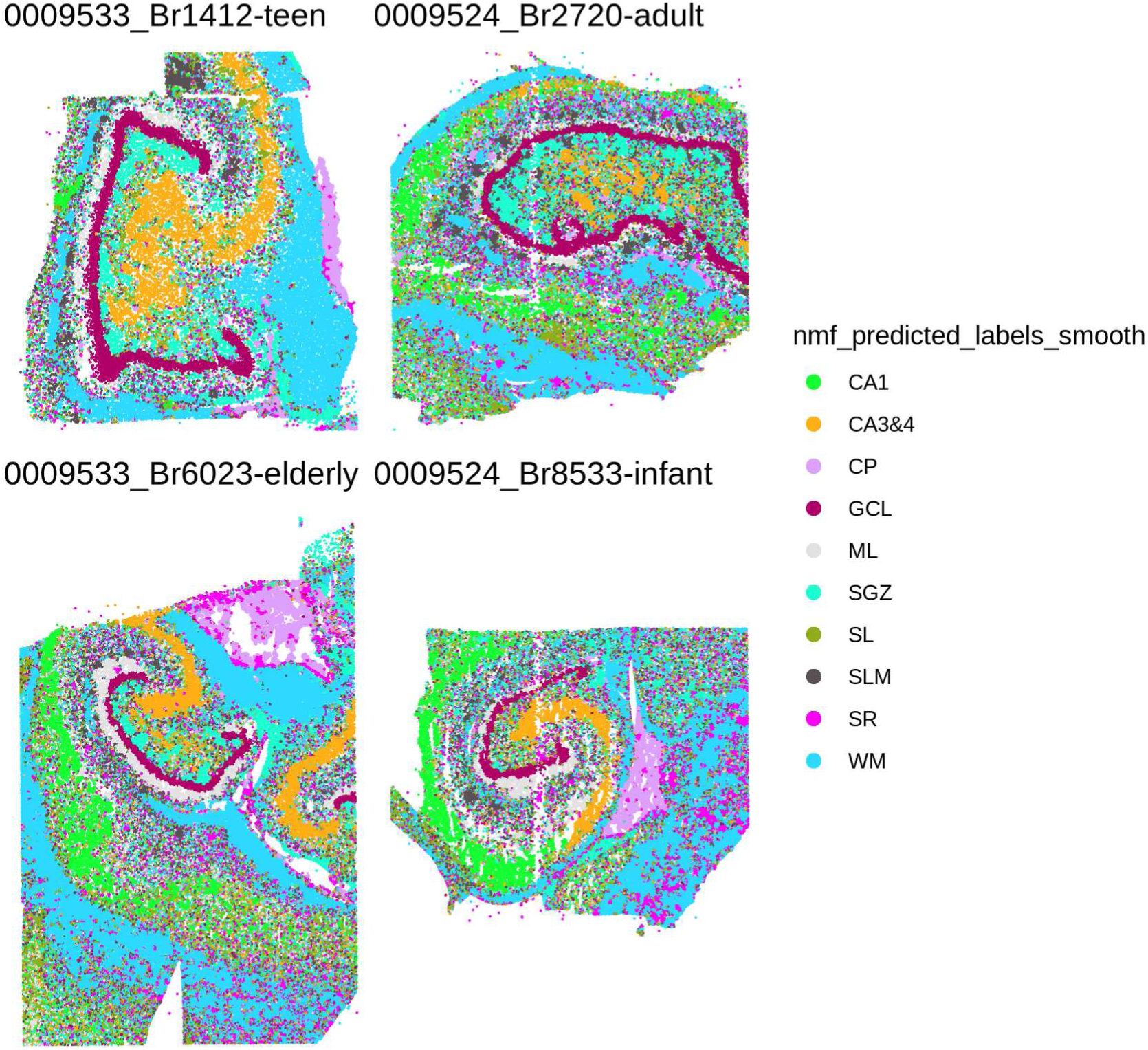
Spotplots of predicted spatial domain labels from spaTransfer (smoothed). Similar to S12, but smoothing (see Methods) has been applied to the raw predicted labels.

**Supplementary Figure S14:**
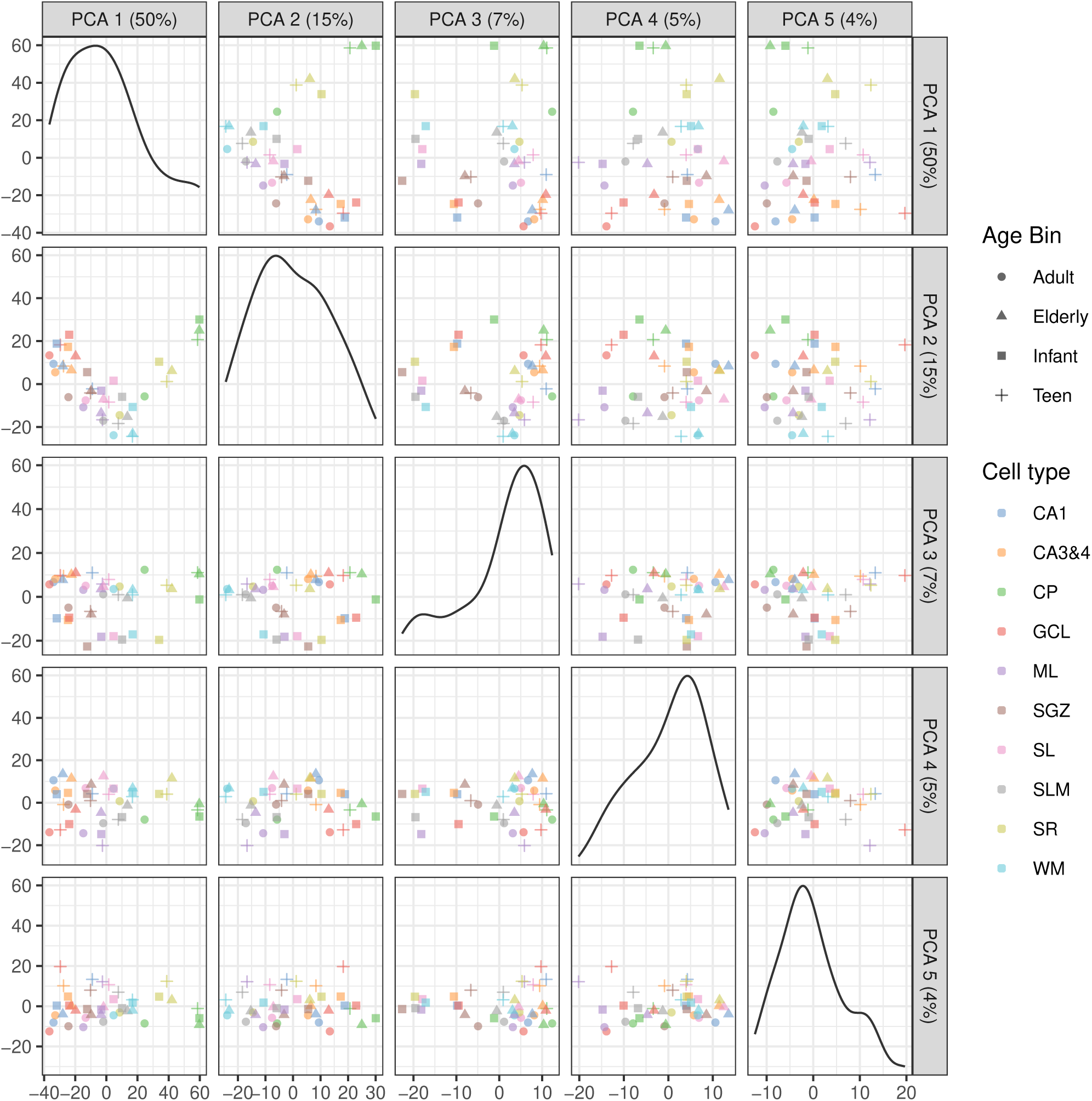
Principal component plots for the Xenium HPC dataset with points colored by the labels transferred from the reference Visium dataset, including CP. The four Xenium HPC samples were pseudobulked to the donor-domain level and PCA was performed on the pseudobulked samples. Each point in this plot represents a unique donor-domain combination. The pseudobulked samples are plotted in PC space PC1 (50% of variance explained) appears to be driven by the CP domain..

**Supplementary Figure S15:**
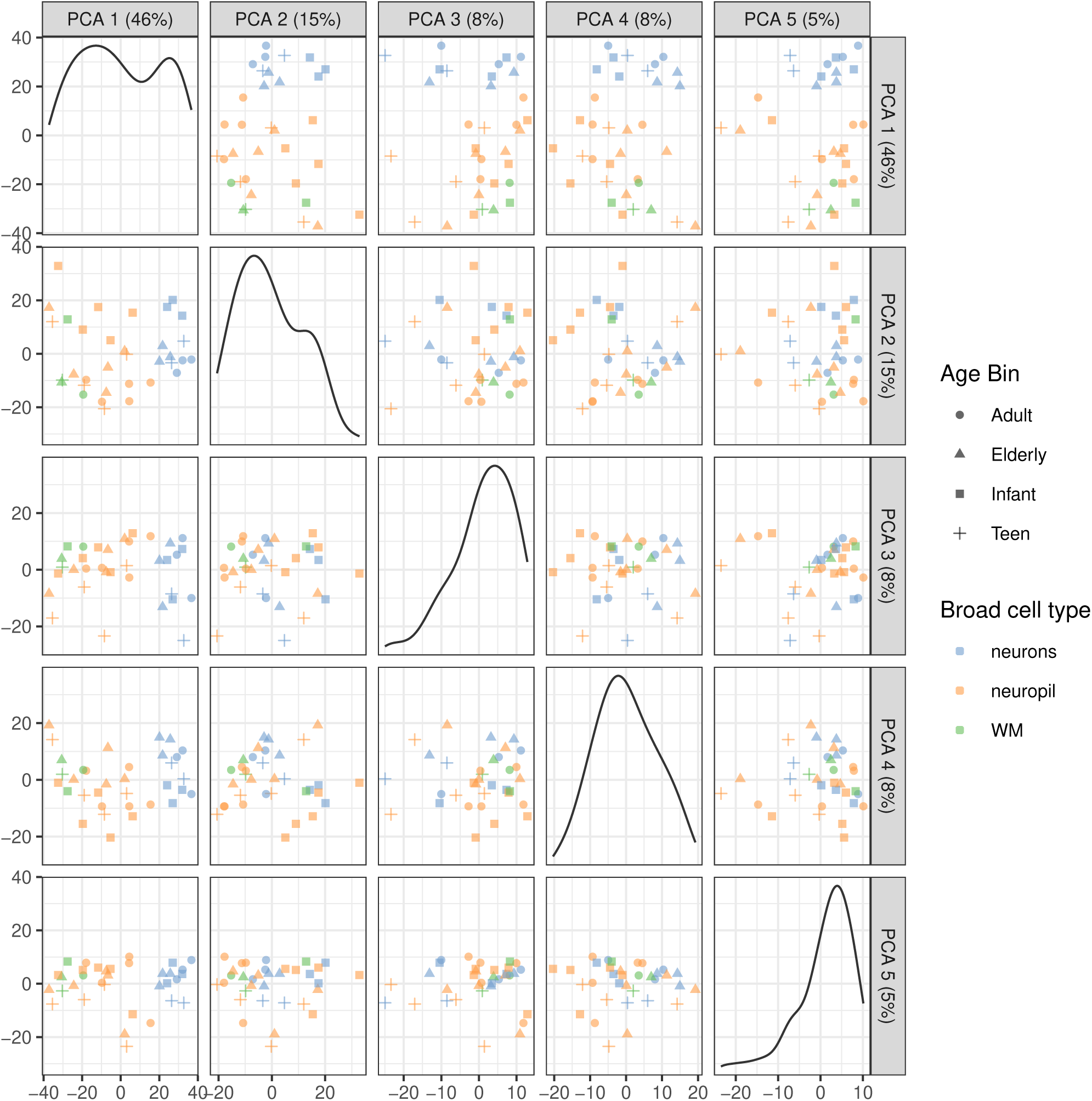
Principal component plots for the Xenium HPC dataset with points colored by the broad cell type of each pseudobulk sample. Same as S14, but with the points colored by broad cell types (neurons, neuropil, white matter). The first principal component (26% variance explained) appears to be driven by the difference between neurons and the other two broad cell types.

**Supplementary Figure S16:**
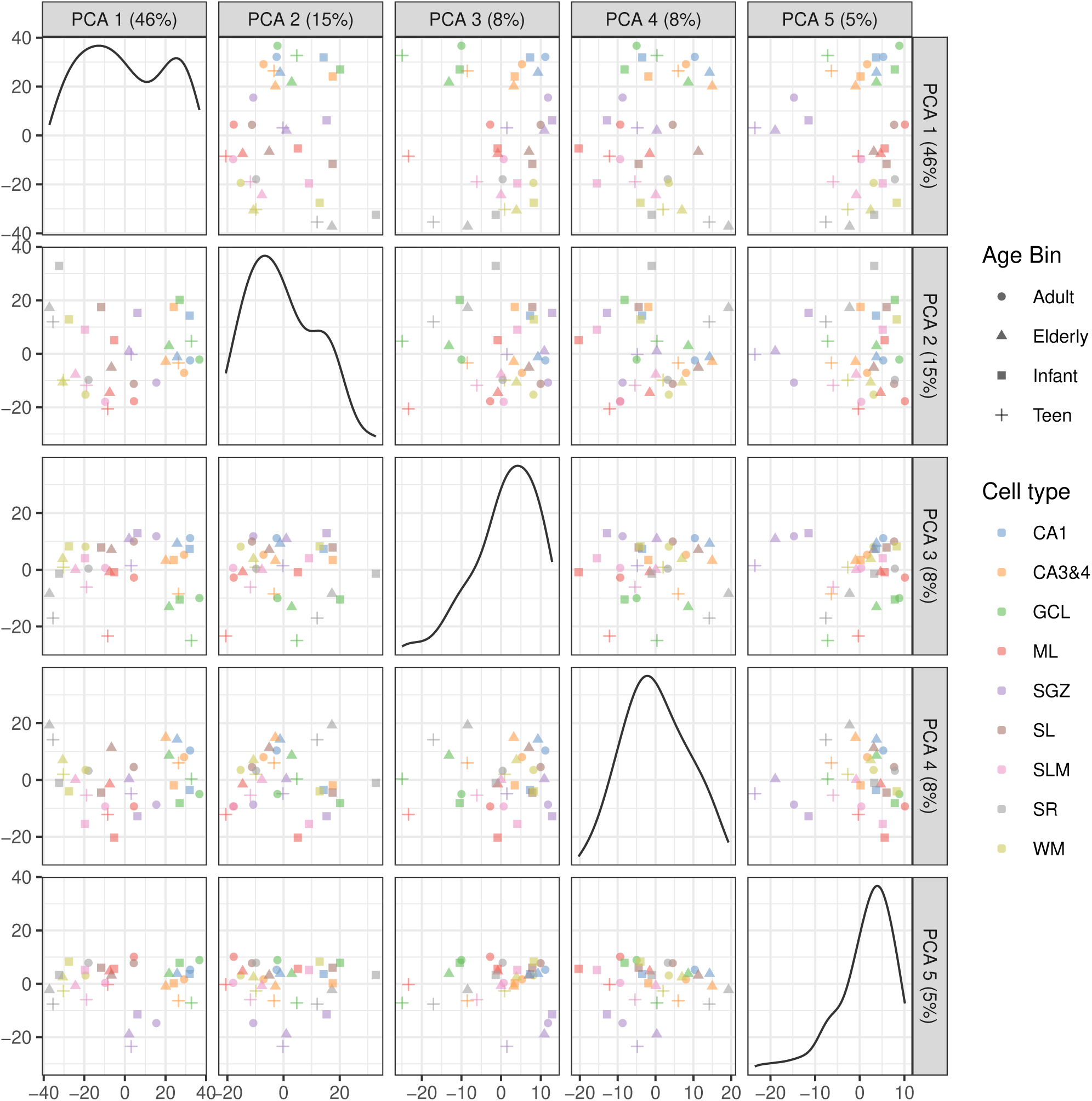
Principal component plots for the Xenium HPC dataset with points colored by the labels transferred from the reference Visium dataset, after removing CP and other donor-specific artifacts. Same as S14, but with the pseudobulk samples representing CP and the teen CA1 and SL regions removed. Principal component 2 (15% of variance explained) appears to be driven by the difference between infant samples and other age groups, suggesting gene expression in the DG can vary across the lifespan.

**Supplementary Figure S17:**
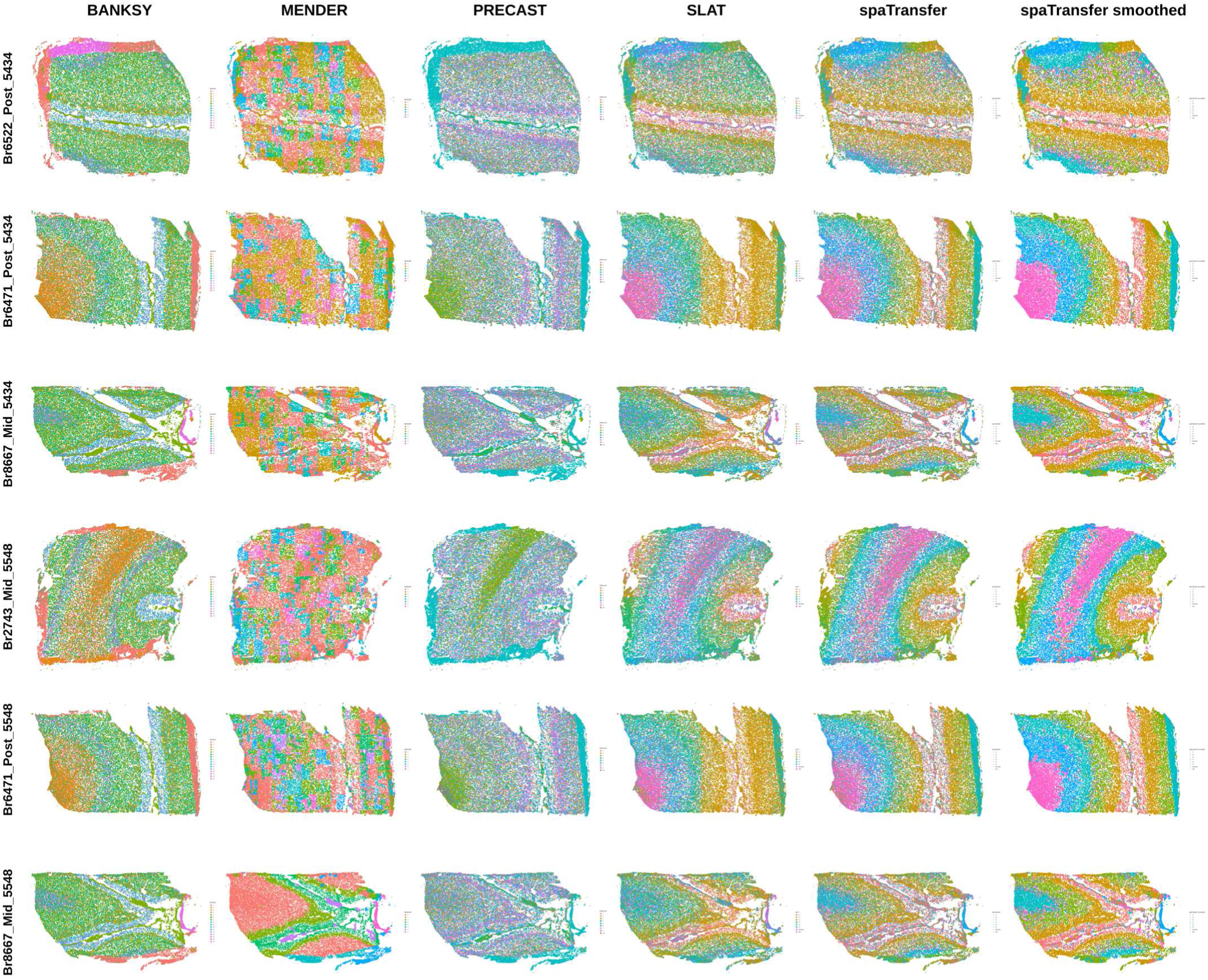
Spotplots of predicted spatial domain labels from each of the label transfer and clustering algorithms applied to the Xenium dlPFC data. Each row represents one of the six tissue sections from the Xenium dlPFC dataset. The columns contain results from (i) BANKSY, (ii) MENDER, (iii) PRECAST, (iv) SLAT, (v) spaTransfer, and (vi) spaTransfer with spatial smoothing.

**Supplementary Figure S18:**
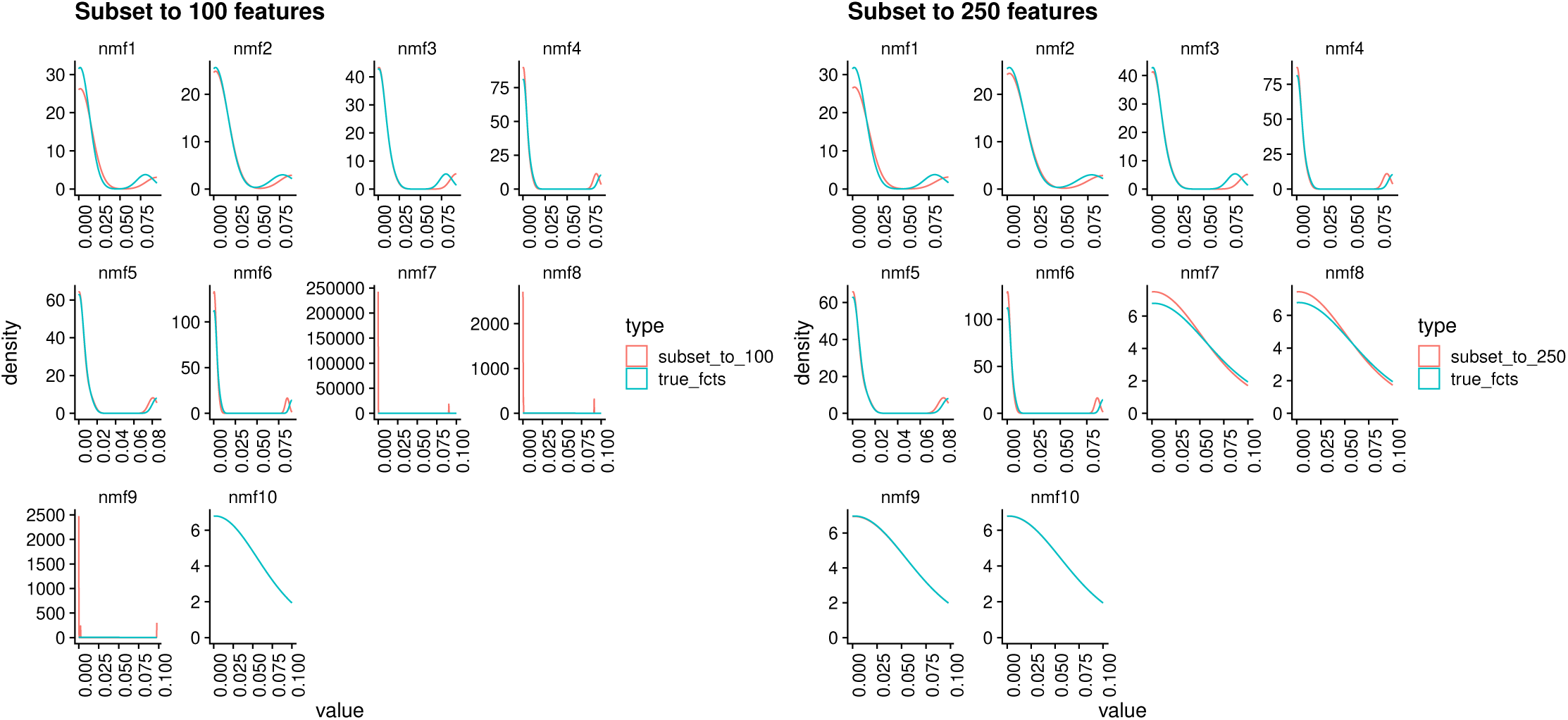
Density plots of learned factors from NMF simulations. A 1000 by 100 matrix was first generated by sampling randomly from a uniform distribution. NMF was performed on this matrix to learn the “true” factors. Then, 100 features (rows) were randomly sampled from the original matrix and a new set of factors were learned by projecting the 100 features with the loadings matrix from NMF. The density of the “true” factors and the factors learned from the 100 features were compared by plotting their densities in blue and red respectively. The same was done with 250 features. Using 100 features results in some factors having different densities compared to the true factors, but at 250 features, the densities of the factors align with that of the true factors.

**Supplementary Figure S19:**
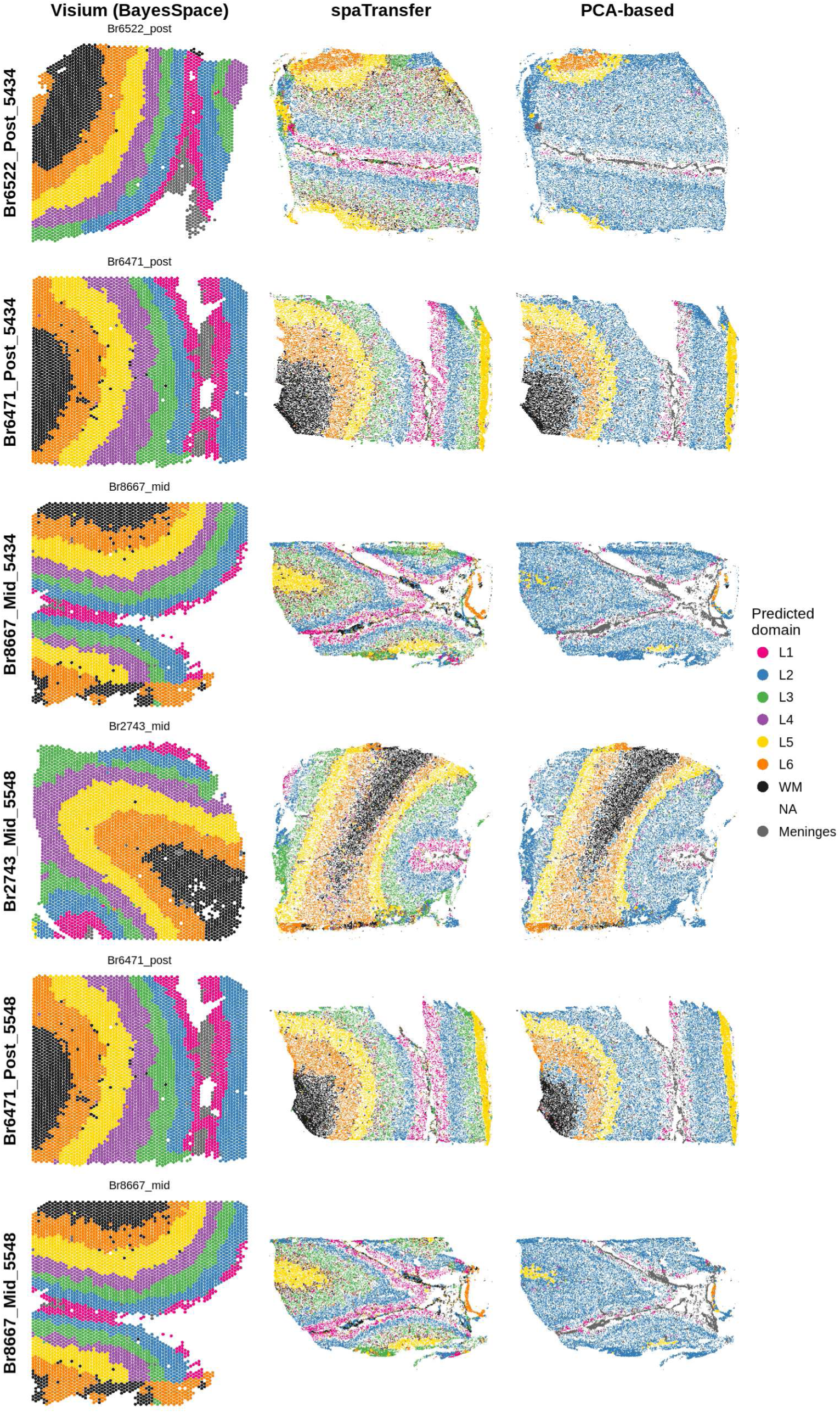
Comparison of NMF- and PCA-based prediction performance when transferring labels from Visium dlPFC data to Xenium dlPFC data. Left column: Visium dlPFC ground truth cortical layer annotations derived from BayesSpace clusters. Middle: spaTransfer NMF-based Xenium cortical layer predictions at rank 25. Right: PCA-based Xenium cortical layer predictions at rank 25. The Visium tissue section in each row is from the same donor and anatomical region as the Xenium section in that row.

